# Charting Postnatal Heart Development Using *In Vivo* Single-Cell Functional Genomics

**DOI:** 10.1101/2025.03.10.642473

**Authors:** Haofei Wang, Yanhan Dong, Yiran Song, Marazzano Colon, Nicholas Yapundich, Shea Ricketts, Xingyan Liu, Gregory Farber, Yunzhe Qian, Li Qian, Jiandong Liu

**Affiliations:** Department of Pathology and Laboratory Medicine, Chapel Hill, NC 27599, USA; McAllister Heart Institute, University of North Carolina, Chapel Hill, NC 27599, USA

**Author notes:** Correspondence: Jiandong Liu, PhD or Li Qian, PhD Department of Pathology and Laboratory Medicine, McAllister Heart Institute, University of North Carolina at Chapel Hill, 111 Mason Farm Road, Chapel Hill, NC 27599, USA. or. These authors contributed equally to this article (Haofei Wang, Yanhan Dong, Yiran Song).

**Keywords:** Cardiomyocyte maturation, Cardiac development, *in vivo* perturb-seq, Spatial transcriptomics, Functional genomics

## Abstract

The transition at birth, marked by increased circulatory demands and rapid growth, necessitates extensive remodeling of the heart’s structure, function, and metabolism. This transformation requires precise spatial and temporal coordination among diverse cardiac cell types; central to this process is cardiomyocyte maturation, yet the regulatory mechanisms driving these changes remain poorly understood. Here, we present a temporal and spatial atlas of postnatal hearts by integrating single-nucleus transcriptomics with image-based spatial transcriptomics, which uncovers the dynamic regulatory networks of cardiomyocyte maturation. To functionally interrogate candidate regulators *in vivo*, we developed Probe-based Indel-detectable Perturb-seq (PIP-seq), a high-throughput platform that uses probe-based chemistry to directly capture sgRNA expression, perturbation status, and transcriptomic profiles at single-nucleus resolution. Applying PIP-seq to postnatal cardiac development identified 21 novel regulators of cardiomyocyte maturation, highlighting critical nodal points in this process. Our study establishes a high-resolution framework for dissecting postnatal heart development, underscoring the integrative and highly ordered roles of microenvironment and intercellular communication in cardiomyocyte maturation. Importantly, PIP-seq enables systematic, high-throughput exploration of gene function and networks underlying complex biological processes in their native *in vivo* context.

## Introduction

A central question in organ development is how the spatial organization and coordinated interactions of different cell types are regulated to form complex functional structures. A critical aspect of this process is the emergence of distinct cell states with specialized activities and niches that cooperate to generate and maintain these larger structures. (*1*, *2*) Despite its significance, the interactions among these components and their contributions to the overall structural and functional integration remain incompletely understood. Consequently, it is imperative to elucidate the components of these cellular ecosystems and uncover the underlying principles that govern their internal organization and interactions(*3–8*). Postnatal cardiac development represents a critical phase during which the heart transitions from a relatively immature organ to a fully functional structure capable of sustaining the circulatory demands of an organism. This process involves coordinated changes in epigenetic, cellular composition, tissue architecture, and mechanical properties that collectively enhance the heart’s efficiency and resilience(*9–13*). A key event during this period is the maturation of cardiomyocytes (CMs), which transition from a fetal state characterized by high proliferative potential and immature contractile machinery to an adult phenotype marked by specialized structural and functional attributes (*14–17*). This includes the development of organized sarcomeres, increased mitochondrial density, and metabolic shifts to oxidative phosphorylation, all of which contribute to the enhanced contractile force and endurance of the adult heart. Understanding the mechanisms that regulate CM maturation and their integration with broader tissue-level changes remains a significant challenge with important implications for both developmental biology and regenerative medicine.

Despite substantial advancements, the study of postnatal cardiac development has predominantly relied on approaches that, while insightful, are limited in their ability to fully capture the intricate mechanisms regulating CM maturation. Recent studies using single-cell RNA sequencing (scRNA-seq) have offered detailed transcriptomic profiles(*18–24*); however, these approaches inherently lose critical spatial information due to tissue dissociation. Moreover, they frequently lack robust functional validation, typically conducted using low-throughput *in vivo* knockout or knockdown strategies. Such traditional methods are constrained by inefficiency, limited resolution, and the risk of secondary effects caused by global gene disruption. While *in vitro* functional genomics platforms offer high-throughput capabilities, their inability to mimic the complexity and physiological relevance of the *in vivo* environment limits their utility in accurately modeling CM maturation(*25*, *26*). The rapid development of CRISPR/Cas9 genome editing tools and single-cell genomic approaches has enabled the establishment of *in vivo* Perturb-seq, a functional genomic platform allowing simultaneous genetic perturbation and functional readout of cells within their native environment(*27–30*). However, even *in vivo* Perturb-seq inherits some disadvantages associated with single-cell-based RNA-seq, including biases introduced by stress-induced transcriptional artifacts during single-cell dissociation(*31*) and low efficiency in capturing cell types that are large or difficult to dissociate. These limitations have restricted the application of *in vivo* Perturb-seq in studying postnatal hearts.

In this study, we constructed a comprehensive postnatal heart atlas by integrating single-nucleus transcriptomic profiling with image-based spatial transcriptomics. This integrative approach captured the spatial and temporal dynamics of postnatal heart development, revealing critical regulatory mechanisms and identifying potential regulators of CM maturation. To functionally investigate these candidates in a high-throughput and *in vivo* context, we developed Probe-based Indel-detectable Perturb-seq (PIP-seq), a platform that uses probe hybridization-based chemistry to simultaneously detect sgRNA and transcriptomic profiles from fixed single nuclei, providing an integrated method to identify perturbation states and their transcriptional consequences. In a proof-of-principle screen, we utilized PIP-seq to analyze 53 candidate regulators of CM maturation, identifying 21 novel regulators and providing functional evidence for their roles in this process. Overall, this study establishes a comprehensive framework for dissecting the mechanisms underlying postnatal heart development by integrating transcriptomic and spatial analyses with functional genomics, providing a rigorous platform for investigating gene function, regulatory networks, and cellular interactions within their native physiological context.

## Result

### Postnatal heart atlas with spatial and temporal resolution

We integrated single-nucleus RNA profiling with single-cell in situ spatial transcriptomic mapping to establish a high-resolution spatial and temporal atlas of the postnatal hearts. Whole hearts were collected at postnatal Day 0 (p0), p7, p14, and p21, capturing the developmental transition from the hyperplastic to the maturational hypertrophic phases of postnatal heart development (Fig. 1A-C). To address the challenge of segmenting multinucleated CMs, we incorporated multimodal cell segmentation staining into our Xenium workflow (Fig. 1B and Fig.S1A). Following stringent filtering of low-quality cells and transcripts, the dataset comprised 67,000 nuclei from snRNA-seq profiling and 402,000 cells from Xenium analysis (Fig.1D and fig.S1B). The combined snRNA-seq and single-cell spatial imaging datasets demonstrated a strong Spearman correlation in gene expression, confirming the high quality of both datasets (Fig.1E and fig.S1C). Subsequently, we constructed a single-cell atlas of the postnatal heart by integrating snRNA-seq datasets with Xenium datasets using cellular Spatial Positioning Analysis via Constrained Expression alignment (CytoSPACE) (*32*)(Fig.1F-G), providing a comprehensive and high-resolution spatial characterization of postnatal heart development across key stages of the maturation process.

**Figure 1.**
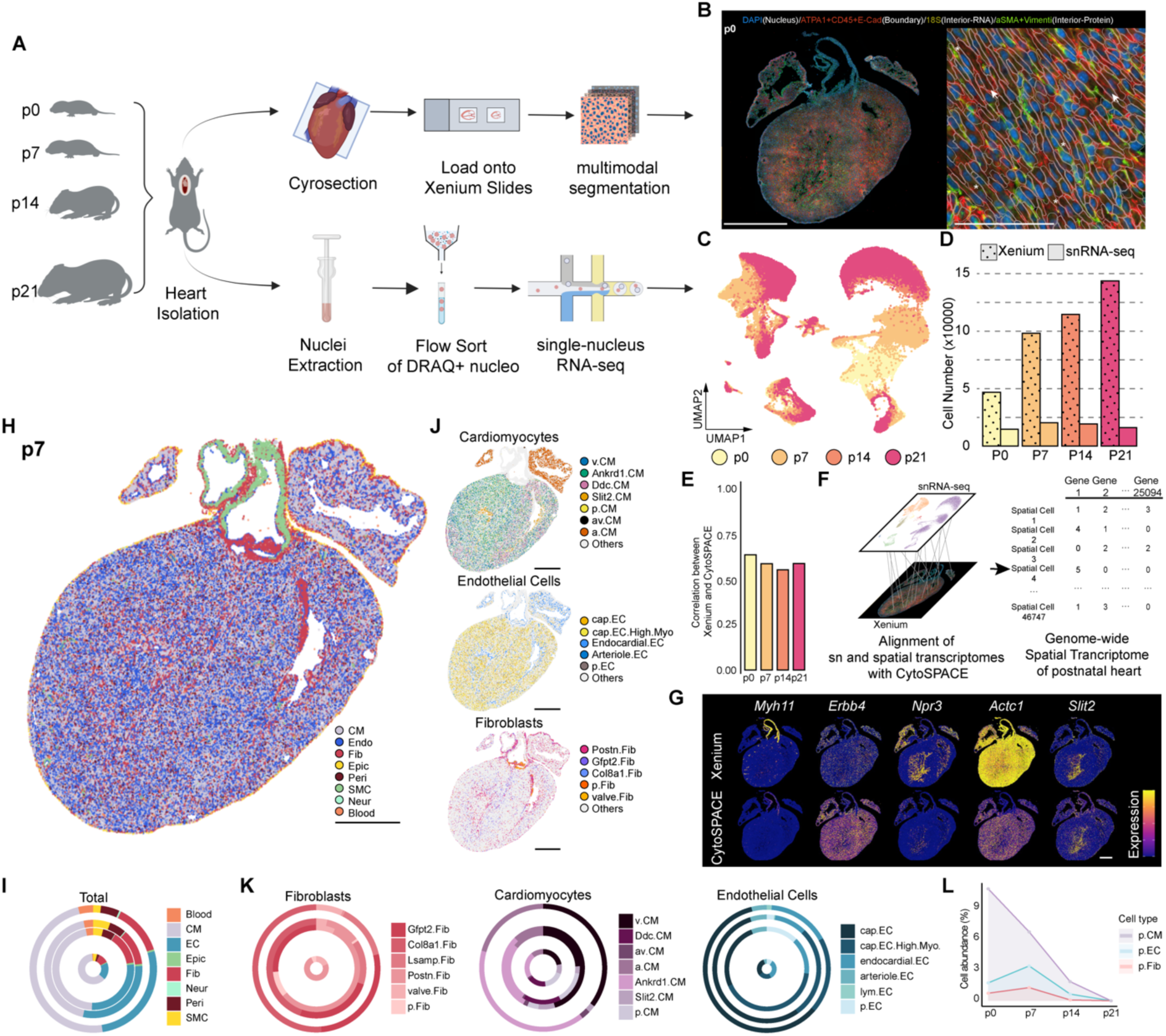
Postnatal heart atlas with spatial and temporal resolution. A) Postnatal heart atlas was composed of two sets of paired data (Single-cell Spatial Transcriptome: Xenium; Single-nucleus Transcriptome: 10x 3’ Gene Expression Library) collected at p0, p7, p14, and p21 of the postnatal development. B) Multimodal segmentation enables improved segmentation of the multinucleated cardiomyocyte (indicated by the asterisk) and enucleated cardiomyocytes as artifacts of section thickness (indicated by the arrow). C) Approximately 67,000 cardiac cell nuclei were clustered into distinct cell populations as shown in the UMAP and across four postnatal timepoints (colored accordingly) D) The number of cardiac cells profiled for Xenium and single-nucleus RNA-seq at each time point. E) Pearson correlation value between the single-nucleus RNA-seq and Xenium for each time point. F) High-resolution alignment of single-nucleus and spatial transcriptomes with CytoSPACE led to the genome-wide spatial transcriptome of postnatal heart development. G) Transcriptome alignment was validated by comparing the normalized expression pattern of selected markers between the original Xenium transcriptome and the CytoSPACE imputed transcriptome. Scale bar, 500μm. H) Spatial map of heart at postnatal day 7. Each cell type was labeled with different color. I) Donut plot showing the changes in cell proportion over the postnatal development. The total cell number of each time point is correlated with the radius of the donut. J) Spatial map showing the subordinate cell states of cardiomyocytes (top), endothelial cells (mid), and fibroblasts (bottom) in the heart at postnatal day 7. K) Donut plot showing the changes in the proportion of the subordinate cell states of fibroblasts (left, red), cardiomyocytes (mid, purple), and endothelial cells (right, blue) over the postnatal development. The total cell number of each time point is correlated with the radius of the donut. L) Line plot showing the abundance of proliferative CMs, proliferative ECs, and proliferative fibroblasts in the heart throughout the postnatal development. The Y-axis represents the percentage of each cell type in the heart. The X-axis represents the time points in postnatal development.

To annotate the cells in our atlas datasets, we employed a combination of an unbiased clustering approach, IKAP (Identifying K mAjor cell Population groups)(*33*), and knowledge-based cell annotation. Our postnatal heart atlas recovered nine major cell types (Fig. 1H, fig. S2A), among which CMs, endothelial cells, and fibroblasts emerged as the three most prominent cell types in the postnatal heart (Fig.1I). With single-cell resolution spatial imaging, these cell types were further subclustered to 25 subordinate cell states (fig.S2B). These subordinate cell states, including 7 CM states, 6 endothelial cell states, and 6 fibroblast cell states, exhibit differentially expressed transcription programs and distinct spatial localization in the postnatal heart (Fig.1J-K, fig.S3). For CM states, we observed CMs highly expressing *Ankrd1* located in the ventricle. Compared to other ventricle CMs (v.CM), these Ankrd1.CM preferentially localizes in the left ventricle and near the base of the heart, a pattern that persists throughout the postnatal development (fig.S4A-C). *Ankrd1* has been reported to be associated with both physiological and pathological stress in the heart, such as those occurring during exercise and myocardial infarction(*34*, *35*, *35–37*). Transcriptomic comparison between *Ankrd1*+ CM and the other ventricle CMs in the postnatal heart revealed a significant upregulation of stress response genes, such as *Atf3* (fig.S4D) (*38*), suggesting that *Ankrd1* expression reflects a location-specific response to mechanical strain accompanying the heart growth. We also observed a unique population of capillary endothelial cells (cap.EC.High.Myo) characterized by a high level of myofiber gene expression, as previously reported(*39*). Spatially, these cells are dispersed throughout the postnatal heart, exhibiting minimal difference in distribution compared to other capillary ECs (Fig.1J, fig.S3, and fig.S5).

The majority of these subordinate cellular states show a sustained existence while demonstrating dynamic changes in abundance throughout postnatal development. For instance, the number of capillary ECs (cap.EC) increases drastically during early postnatal development before reaching a plateau at later stages (Fig.1K), suggesting a tightly coordinated temporal regulation of postnatal coronary development. Consistent with the transition from hyperplasia to hypertrophic growth, the postnatal heart atlas also captured transient proliferative cell states in CMs, endothelial cells, and fibroblasts. Unlike other cell states with distinct spatial locations, these proliferative cell states were evenly distributed across both atrial and ventricular chambers (Fig.1J, fig.S6). Interestingly, these distinct proliferative cell states exhibit highly similar temporal dynamics, peaking at p7 and largely diminishing by p21 (Fig. 1L), suggesting tightly coordinated and regulated postnatal maturation across different cardiac cell types. Taken together, the postnatal heart atlas offers a high-resolution, comprehensive blueprint of individual cellular states, defined by distinct transcriptomic profiles and spatial organization in the postnatal heart.

### Localized regulatory mechanism of postnatal heart development

The spatially orchestrated cellular interactions among specialized cardiac cell types are critical for regulating postnatal cardiac development, highlighting the need to characterize stable and distinct cellular neighborhoods (niches) across postnatal stages(*40*). A total of 12 cell niches were identified (Fig.2A, fig.S7), eight of which remain consistently present across all postnatal stages. Notably, these niches align with well-defined anatomical structures in the postnatal heart (Fig.2A, fig.S7). For example, niche 1, characterized by an abundance of endocardial endothelial cells(endo.EC) and trabecular CMs (Slit2.CM), is associated with the trabecular structures of the ventricle (Fig.2A, fig.S7,S8). Niches 3 and 8 map to the atrium of the postnatal heart, while the Ankrd1.CM-enriched niche 4 is consistently situated in a peri-arteriole location within the ventricular region of the postnatal heart (Fig.2A, fig.S7,S8). Interestingly, all four timepoint-specific niches were confined to the ventricular chamber, highlighting the dynamic nature of ventricular remodeling during postnatal development. Niche 0, established by p7 and expanding thereafter, is enriched with pericytes (Peri) and capillary ECs (cap.EC), reflecting a continuous expansion of the coronary plexus in the postnatal heart (Fig.2B, 2C, fig.S7). Niche 2 (p7), 9 (p0), and 11 (p14, p21) are mostly enriched with ventricular CMs at their respective time point. Notably, the early-stage specific niches 2 and 9 contain more proliferative CMs than the late-stage niche 11, highlighting a decline in regenerative capacity during postnatal heart development (Fig.2A, fig.S7,S8).

**Figure 2.**
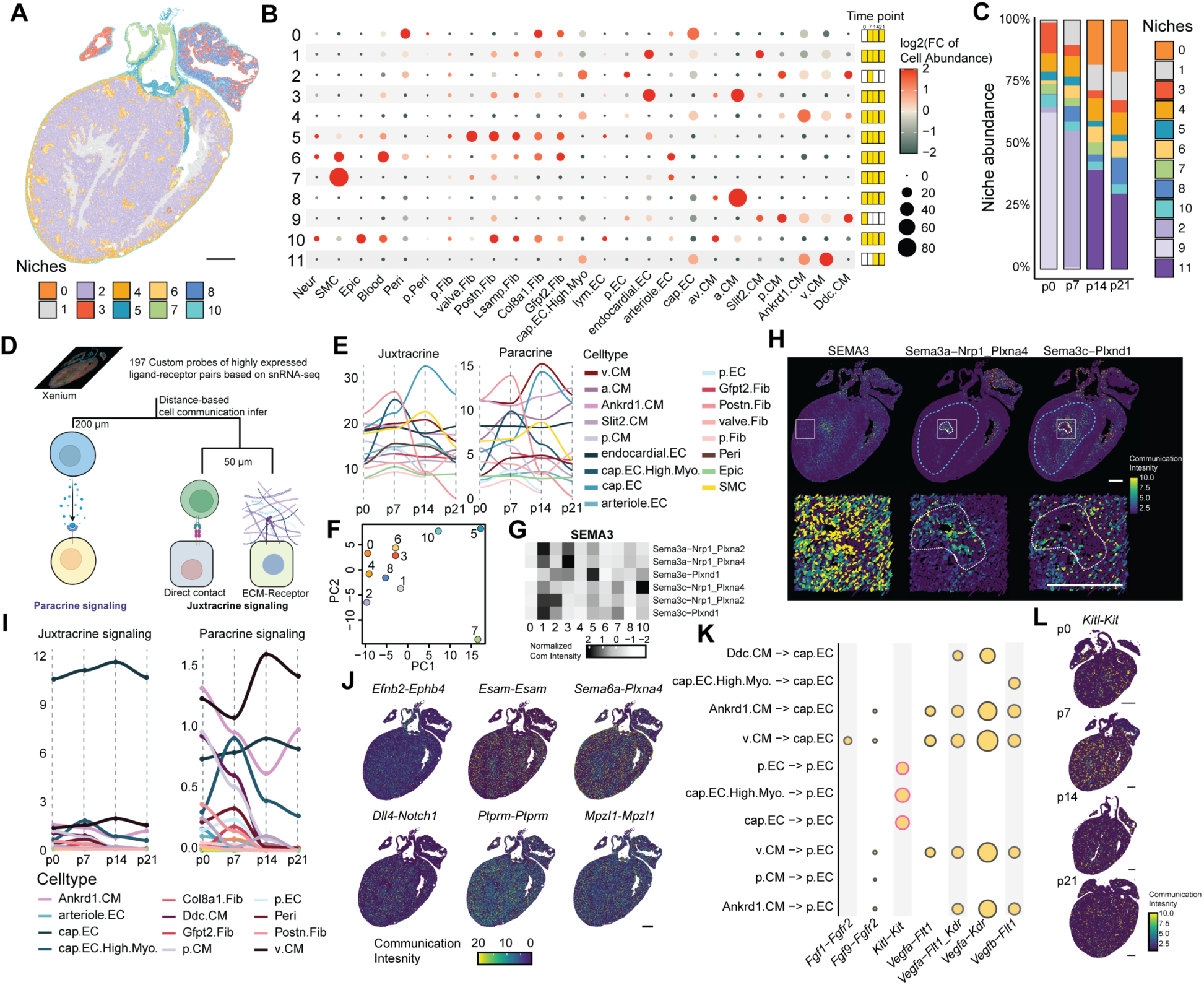
Localized regulatory mechanism of postnatal heart development. A) Spatial map of heart at postnatal day 7. Each niche was labeled with different colors. Scale bar, 500μm. B) Postnatal heart was categorized into 11 niches with distinct cellular compositions. Cell type abundance was presented by the size and color of corresponding dot. For each niche, the time window it presented in the postnatal heart is highlighted as yellow boxes. C) Stacked bar plot showing the proportion of each niche in the postnatal heart throughout the postnatal development. D) Distance-based inference of cell communications using Xenium data. Penalty was applied to the paracrine signaling if the cell pair has a distance above 200 μm and to juxtracrine signaling if the cell pair has a distance above 50 μm. E) Line plot showing the juxtracrine (Left) and paracrine (Right) signaling intensity sent by each cell type throughout the postnatal development. The Y-axis represents the cumulated communication intensity sent by each cell type. The X-axis represents the time point in the postnatal heart. F) PCA plot showing the overall cellular communication difference between niches in the postnatal heart at day 7. G) Heatmap showing the relative communication intensity of each ligand-receptor pair of the SEMA3 pathway at different cellular neighborhoods of the postnatal day 7 heart. H) The spatially resolved activities of the SEMA3 pathway and two ligand-receptor pairs, Sema3a-Nrp1_Plxna4 and Sema3c-Plxnd1 at postnatal day 7 heart. The white dashed area highlights the differential spatial activities of two ligand-receptor pairs. Scale bar, 500μm. I) Line plot showing the juxtracrine (Left) and paracrine (Right) communication intensity received by capillary ECs in the ventricle niches throughout the postnatal development. The Y-axis represents the cumulated communication intensity received by cap.ECs. The X-axis represents the time point in the postnatal heart. J) The spatially resolved activities of *Efnb2*-*Ephb4*, *Esam*-*Esam*, *Sema6a*-*Plxna4*, *Dll4*-*Notch1*, *Ptprm*-*Ptprm*, and *Mpzl1*-*Mpzl1* at postnatal day 7 heart. The color Scheme represents the normalized communication intensity. Scale bar, 500μm. K) Dot plot showing the activated paracrine ligand-receptor pairs received by the cap.EC and p.EC. The p.EC specific Ligand-receptor pairs were highlighted with red border. The size of each dot represents the normalized communication intensity of each ligand-receptor pair. L) The spatially resolved activities of *Kitl*-*Kit* throughout the postnatal development. The color Scheme represents the normalized communication intensity. Scale bar, 500μm.

To define the role of these niches during the structure and function maturation of postnatal heart development, we first set out to infer the activated cellular communications within the postnatal heart. Using 197 custom probes targeting highly expressed ligand-receptor pairs identified from our snRNA-seq, we performed a spatial-resolved cell communication analysis to construct the cellular communications network (Table.S1). Specifically, a penalty is incorporated to reduce the inferred cell communication probability when the sender and receiver cells exceed a defined distance threshold (Fig.2D)(*41*). Overall, Postn.Fib serve as the primary communication hub via juxtracrine and paracrine signaling during the early stage (p0–p7), while cap.ECs assume this role at p14, signaling a dynamic reorganization of the cellular communication network that orchestrates cardiac maturation. (Fig. 2E, fig.S9A).

To further dissect the cell communication networks within each niche, we next focused on analyzing their distinct signaling dynamics. Principle component analysis of the cell communication activities within each niche revealed an extensive signaling diversity among niches for both paracrine and juxtracrine signaling (Fig.2F). Unsupervised clustering of ligand-receptor pairs captured both the shared and unique Ligand-Receptor (L-R) pairs to each niche (fig.S9B-C). *Sema3c-Plxnd1* interaction is detected in the trabecular (niche 1) and compact ventricle niche (niche 2), consistent with its role in ventricle wall compaction (Fig.2G and 2H)(*7*). Conversely, *Sema3a*, another member of the SEMA3 family, engages with *Nrp1-Plxna1* and shows localized activity within the trabecular niche (niche 2), supporting its role as a neural chemorepellent in postnatal cardiac sympathetic innervation (Fig.2G and 2H)(*42*). The spatial segregation of these L-R pairs underscores the localized coordination required for postnatal heart development.

We then applied the cellular communication analysis on the ventricle niches to study coronary plexus expansion, a critical event during postnatal heart development. Focusing on the communication received by the cap.EC within the niche, cap.EC itself dominates the rest of cardiac cells to interact with neighboring cap.EC through juxtracrine signaling, while the v.CM and Ankrd1.CM functions as a major sender to the cap.EC through paracrine signaling (Fig.2I). Further analysis of the active juxtracrine L-R pairs received by cap.EC not only confirmed several well-established L-R pairs implicated in capillary angiogenesis, including *Efnb2-Ephb4*(*43*), *Esam- Esam*(*44*), *Sema6a-Plxna4*(*45*), *Dll4-Notch1*(*46–48*), but also identified previously uncharacterized L-R interaction (Figure 2J, fig.S11). Among these, *Mpzl1*, a component of the cell surface receptor protein tyrosine kinase signaling pathway, was found to be activated among the cap.ECs (Fig.2J, fig.S11). For paracrine signaling, beyond the well-known VEGF signaling pathway(*49*), our L-R pair analysis revealed that *Fgf1-Fgfr2*, from v.CM to the cap.EC, also played a pivotal role in the angiogenesis in the postnatal heart (Fig.2K). Next, we examined the L-R pairs received by the proliferative ECs in the postnatal heart, as these proliferative ECs are the stalk cells responsible for the elongation of sprouting capillary branches. The *Kitl-Kit* L-R pair emerged as one of the dominant and specific paracrine communications received by the proliferative ECs, with its peak activities at p7 (Fig.2K and 2L). Given its reported role in cellular migration(*50*), the activation of the *Kitl-Kit* signaling axis could also be involved in endothelial cell migration to support the capillary vessel sprouting.

Building on the intricate cell communications that orchestrate postnatal heart development, it is equally critical to understand how these pathways converge on specific transcription factors (TFs) to refine and sustain the gene regulatory networks underlying heart maturation. Taking advantage of the genome-wide spatial expression in our heart atlas, we performed single-cell regulatory network inference and clustering (SCENIC)(*51*) analysis to identify activated gene regulatory networks (regulons) in the postnatal heart. Visualizing regulon activities using UMAP revealed that most non-myocyte populations exhibit gradual and progressive variation in regulon activities across postnatal stages. In contrast, CMs form two clusters corresponding to p0-p7 and p14-p21 time points, indicating substantial transcription network rewiring at p7 (Fig.3A). To evaluate the temporal dynamics of inferred regulon activity, we performed semi-supervised clustering of regulon activity, incorporating both cell type annotations and temporal information (Fig.3B, fig.S12). In most non-myocytes, regulon activities remain largely stable throughout postnatal maturation, whereas CMs exhibit distinct sets of regulons with differential activity during this period (Cluster 5, 11, and 12). In addition to identifying well-characterized transcription factors driving the cell type-specific transcription programs, the SCENIC analysis also uncovers TFs with previously unreported roles in postnatal heart development (Fig.3B, fig.S13,S14,S15). For instance, *Fosl2*, an AP-1 transcription factor, has not been directly implicated in postnatal heart development. However, based on our analysis, it exhibited specific activity in the epicardium of postnatal heart (fig.S13, cluster 4). In line with the spatially diversified cell communications, the regulon activity also exhibited a wide range of Moran’I scores representing clustered, dispersed, or random patterns in the postnatal heart(*52*). Of note, *Rfx2*, topping the 200 regulons, showed the highest Moran’I score with highly clustered activities in the atria (Fig.3C and 2D). Interestingly, *Foxc1* is another top regulon with a high Moran’I score. It showed a distinct activity in the inner lining of both the cardiac chamber and outflow tract, consistent with its reported role in the morphogenesis of the cardiac outflow tract and ventricular chamber (Fig.3C and 3D)(*53–55*).

**Figure 3.**
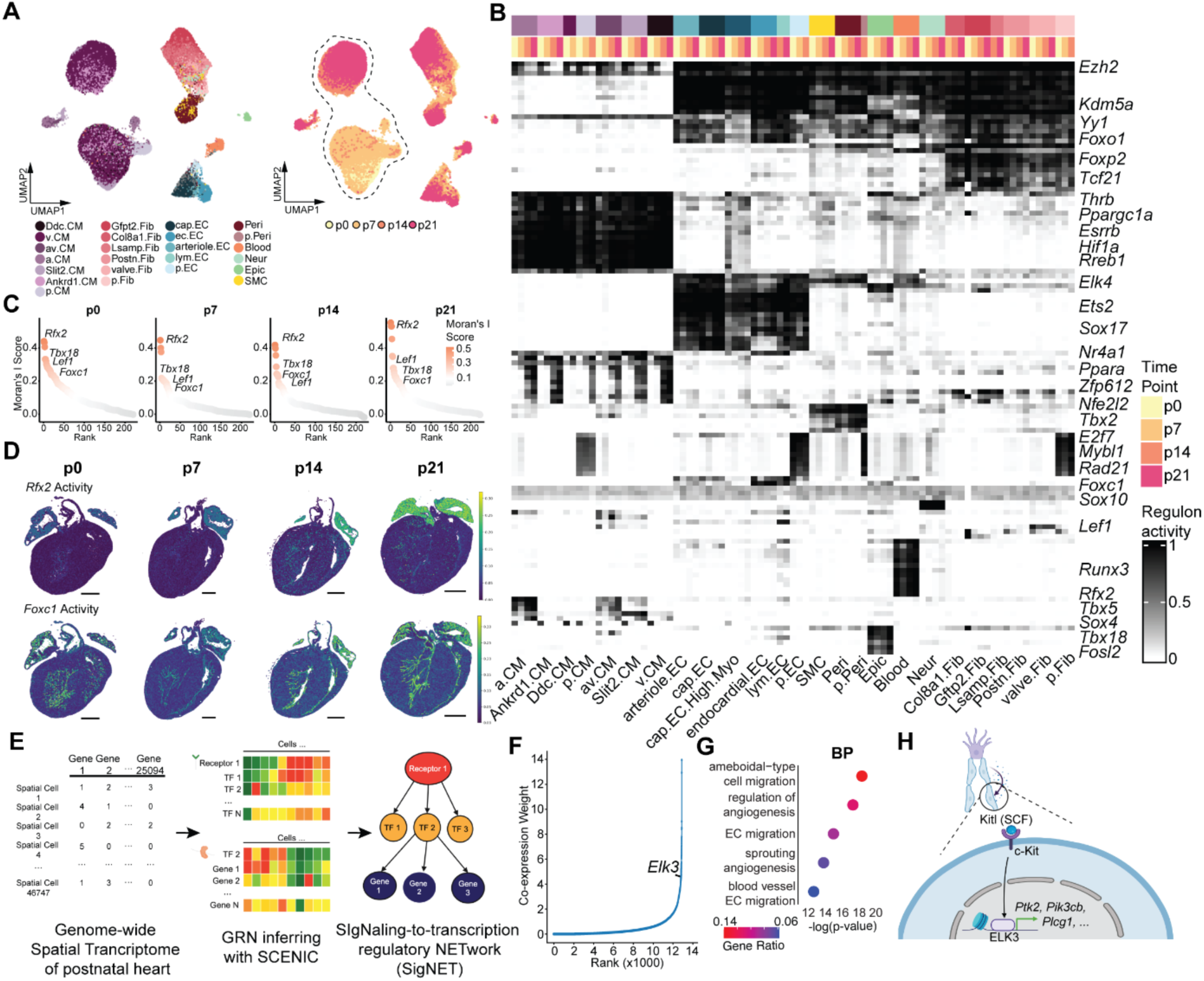
SigNET reconstructed the link between upstream signaling pathways to the downstream transcription factors in the postnatal heart. A) The significant reconfiguration of the regulatory landscape between p7 and p14 cardiomyocytes is evident by the large gap between the CM clusters in regulon activity based UMAP. B) Heatmap showing the regulons with highest activity (top 20) in each cell type at each postnatal time point. The color Scheme represents the regulon activity from high (Black) to low (White). C) Scatter plot showing the Moran’s I Score for each regulon during the postnatal development. D) Regulon activities of *Rfx2* and *Foxc1* were displayed in the postnatal hearts spatially and temporally. The color scheme represents the regulon activity of each cell in the postnatal heart. Scale bar (p0, p7), 500μm; Scale bar(p14, p21), 1000μm. E) SigNET analysis unveils the potential downstream transcription factors of the activated signaling pathways. F) Scatter plot showing the *Elk3* as one of the downstream transcription factors activated by *Kitl*-*Kit* in the postnatal day 7 heart. G) Enriched Gene ontology of the downstream genes controlled by the *Elk3* in the postnatal heart. H) Schematic plot showing that receiving KITL from the neighboring capillary ECs could activate the proliferation program in the proliferative ECs through *Elk3* to support the angiogenesis in the postnatal heart.

Next, to link the upstream signaling pathways to the downstream transcription factors, we established SigNET, Signaling-to-transcription regulatory NETwork, pipeline by uncovering the co-expression patterns among the receptors and its putative downstream transcription factors and then its’ corresponding gene regulatory network in the SCENIC analysis (Fig.3E). With the SigNET, we could reconstruct the regulatory axis from the signaling pathways to the downstream TFs and its target gene programs in the postnatal heart. Applying the SigNET on the *Kitl-Kit* signaling received by the proliferative ECs in the postnatal heart revealed a co-activation of the transcription factor, *Elk3*, in proliferative ECs (Fig.3F). GO enrichment analysis of the gene programs downstream of *Elk3* reveals a significant enrichment of biological processes related to endothelial cell migration and sprouting angiogenesis (Fig.3G). This suggests that during the expansion of the postnatal capillary plexus, the cell migration program is activated in the proliferative ECs through *Elk3* in response to the activation of *Kitl-Kit* from neighboring capillary endothelial cells (Fig.3H).

Altogether, the integrative spatial analyses of the postnatal heart atlas revealed the coordinated effort of the cardiac cells in shaping the postnatal heart, including the capillary angiogenesis and cardiac sympathetic innervation, through spatially restricted ligand-receptor interactions. Simultaneously, we constructed a comprehensive regulatory network by connecting dynamic shifts in transcriptional regulatory networks to extracellular communications through the novel SigNET pipeline. This highlights how factors such as *Elk3* mediate the endothelial migration program in response to Kit activation.

### Spatial insights into cardiomyocyte maturation

Among all cardiac cell types in the postnatal heart, CMs exhibited the highest responsiveness to the postnatal maturation process, as evidenced by the highest Area Under Curve (AUC) scores using a machine-learning framework, Augur (Fig.4A)(*56*). The maturation of CMs is a critical component of postnatal cardiac development, driving structural, metabolic, and functional adaptations to meet increasing physiological demands. Despite its complexity, current studies often rely on limited molecular markers to infer CM maturation status. To establish a robust temporally and spatially resolved measure of CM maturation, we designed probes to detect transcripts of genes associated with maturation programs, including those encoding transcription factors, sarcomere proteins, metabolic enzymes, and electrophysiological regulators within our spatial transcriptomics framework (Fig.4B, Table.S1). We then computed a quantitative maturation index by averaging the standardized expression values of these genes. Mapping this maturation index to the spatial transcriptomics landscape revealed a steady increase in CM maturation from p0 to p21 (Fig.4C and 4D) and, upon closer examination, significant spatial heterogeneity in CM maturation (Fig.4C).

**Figure 4.**
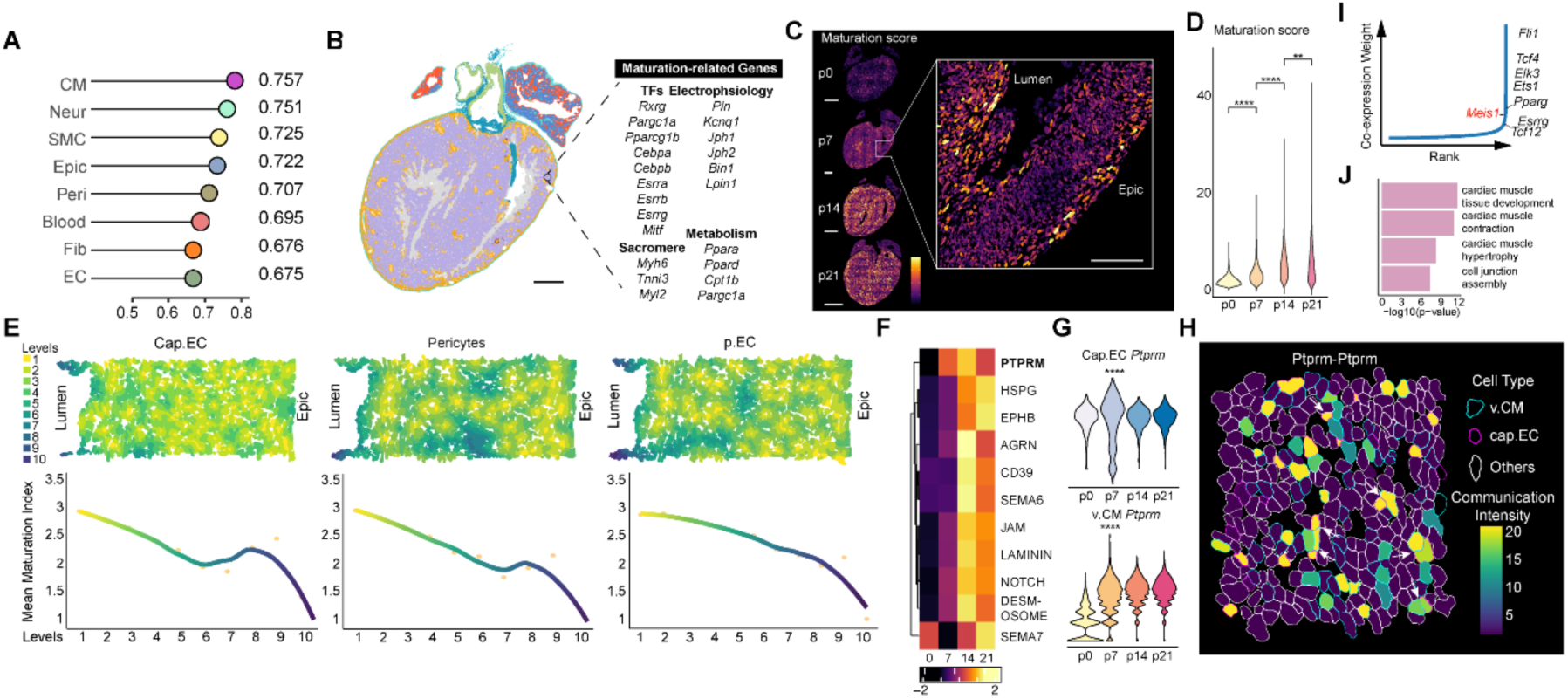
Spatial insights into cardiomyocyte maturation. A) Cell type priority score of each cell type during postnatal development based on Augur analysis. x-axis is the Area Under Curve (AUC) score. B) Maturation Index indicates the cardiomyocyte maturation status through normalized expression of key maturation-related genes. Scale bar, 500μm. C) Spatially mapped maturation score of each cardiomyocyte during postnatal development. Scale bar (p0, p7), 500μm; Scale bar (p14, p21), 1000μm; Scale bar (Zoomed-In view), 250μm. D) Violin plot showing a significantly increased maturation score throughout postnatal development. ****, p<0.00001; **, p<0.001. E) The correlation of maturation index and the cardiomyocyte’s distance to other non-myocyte cell types (Left: Capillary EC, Middle: Pericytes, Right: proliferative EC). Top panel present an example view of the proximity level of neighbouring cells to each cell type. F) Heatmap showing the communication activities of the signaling pathway between the capillary EC and the ventricle cardiomyocyte during postnatal development. G) Violin plot showing the expression of *Ptprm* in capillary EC (Top) and ventricle CM (Bottom) throughout the postnatal development. H) A zoomed-in view of spatially resolved *Ptprm*-*Ptprm* activities in the ventricle of postnatal day 7. Color Scheme represent the cell communication activity from high (yellow) to low (dark blue). I) Scatter plot showing the *Meis1* as one of the downstream transcription factors activated by *Ptprm*-*Ptprm* in the postnatal day 7 heart. J) Enriched Gene ontology of the downstream genes controlled by the *Meis1* in the postnatal heart.

To dissect this spatial heterogeneity, we stratified CMs based on their spatial proximity to non-myocyte cells (Fig.4E) and analyzed the correlation between these spatial relationships and maturation index. Notably, CMs in closer proximity to capillary-associated cell types—including cap.EC, cap.EC.high.myo, p.EC, and pericytes—exhibit significantly higher maturation index, highlighting a potential role of microvascular interactions in promoting CM maturation (Fig. 4E and Fig. S17). Leveraging our spatial transcriptomic map, we explored cellular communication between cap.ECs and CMs. Among the 11 juxtracrine signaling pathways, *Ptprm-Ptprm* showed a localized activity in adjacent CMs and capillary ECs from p7 onward, consistent with its expression profile in our snRNA-seq data (Fig. 4F-H). *Ptprm* encodes a transmembrane protein tyrosine phosphatase implicated in cell differentiation and migration(*57*, *58*), and through SigNET analysis, we identified *Meis1* as its downstream mediator.(Fig. 4I). Gene ontology enrichment analysis of Meis1-regulated transcriptional program inferred from SCENIC analysis revealed enrichment for genes involved in CM proliferation regulation (Fig.4J). This finding supports the reported role of *Meis1* in postnatal CM cell cycle exit (*59*) and suggests that it may serve as a critical link between *Ptprm*-mediated signaling and the regulation of CM maturation through cell cycle exit.

### Intrinsic and extrinsic regulation of cardiomyocyte maturation

CM maturation is regulated by a complex interplay of transcriptional regulation and intercellular signaling, yet its mechanisms remain incompletely understood. As a first step toward delineating the intrinsic transcriptional programs governing CM maturation, we performed Weighted Correlation Network Analysis (WGCNA)(*60*). By clustering transcriptionally similar CMs, this analysis identified four distinct temporally regulated gene co-expression modules (Fig.5A and 5B). Modules 1 and 4 comprise gene programs upregulated at p7 and p14, encircled for genes involved in cardiac contraction, membrane potential regulation, and fatty acid transport (Fig. 5C). Module 2 represents a transcriptional program progressively repressed during postnatal CM maturation, characterized by the downregulation of WNT signaling pathway (Fig. 5C). Next, we reconstructed the upstream intrinsic regulatory network orchestrating these gene expression programs during CM maturation. Leveraging the inferred regulon activity profiles from SCENIC analysis of the heart atlas (fig.S12, cluster 5, 11, and 12), we constructed a temporal gene regulatory network comprising 32 CM-specific regulons across postnatal time points (Figure 5D). 7 of the 32 regulons have been previously implicated in postnatal CM maturation, including those involving the TFs *Thrb*(*61*, *62*), *Ppargc1a*(*23*, *63*), and *Esrra*(*64*). We performed gene ontology analysis on the predicted downstream targets of the 25 novel regulons, revealing their involvement in various maturation-related programs. For example, *Tef* is linked to the regulation of lipid metabolism during postnatal maturation, while *Zfp612* is associated with CM electrophysiological maturation (fig.S16).

**Figure 5.**
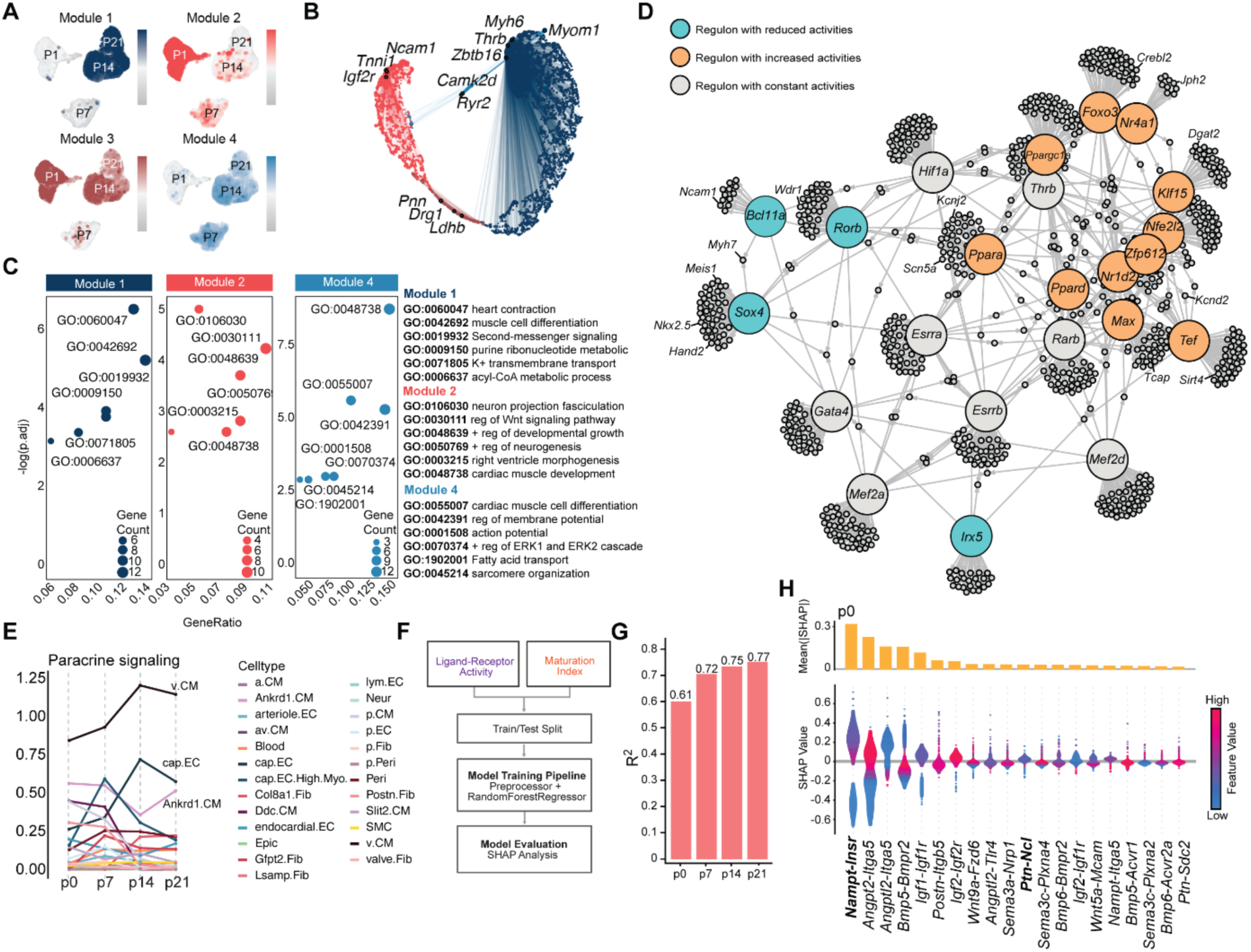
The intrinsic and extrinsic regulation of cardiomyocyte maturation. A) Four modules showing distinct temporal expression behaviors in cardiomyocyte maturation were identified from the WGCNA analysis. B) UMAP plot of the co-expression network in cardiomyocyte maturation. Each node represents a single gene, and edges represent co-expression links between genes and module hub genes. Point size is scaled by kME. Nodes are colored by co-expression module assignment. The top hub genes in each module are labeled. Network edges were downsampled for visual clarity. C) Dot plot showing the significantly enriched gene ontology in each WGCNA module. The size of each dot represents the counts of genes related to each gene ontology. The annotation of each gene ontology is listed at the right of the dot plot. D) Intrinsic regulatory network involved in postnatal cardiomyocyte maturation. Regulons showing different activity patterns were labeled by different colored dots. Blue, regulons with reduced activities during cardiomyocyte maturation; Grey, regulons with constant activities during cardiomyocyte maturation; Yellow, regulons with increased activities during cardiomyocyte maturation. Direct targets in each regulon were downsampled for visual clarity. E) Line plot showing the paracrine communication intensity received by ventricle cardiomyocytes from other cardiac cells throughout the postnatal development. F) Schematic of the training machine learning models on L-R pairs impact on maturation index. G) Bar chart showing the R^2^ of model for each timepoint in inferring CM maturation. Y-axis is the R^2^ and X-axis is the timepoint. H) Top: Bar plot showing the average of the absolute value of SHAP value for each L-R pair. Bottom: Violin plot showing the SHAP value distribution of the L-R pairs with top 20 impact on the model output of p0 heart. Color Scheme reflects the L-R activity for each cell with red as high activity and blue as low activity.

To explore extrinsic signals in CM maturation, we analyzed signaling pathways received by the ventricle CMs. Throughout postnatal maturation, v.CM is the primary sender to neighboring v.CM, followed by capillary ECs (Fig.5E, fig.S18A). Unsupervised clustering of communication probabilities revealed distinct temporal dynamics of the signaling pathways across maturation stages (fig.S18B-C). A key signaling transition involves the decline in endocardial-derived IGF2(*65*) and NRG1(*66–69*), both of which promote CM proliferation. This shift coincides temporally with the transition from hyperplastic to hypertrophic CM growth (fig.S18D-E). Beyond these well-established pathways, our analysis identified additional signaling dynamics, including *Pleiotrophin* (PTN), an early maturation cue secreted by Postn.Fib at P0 that gradually declines over time (fig.S18B and 18F). We then applied machine learning to further evaluate the impact of L-R interrelations on CM maturation. Using random forest regression, we trained models on our dataset incorporating L-R activities from each CM with corresponding maturation indices at each time point (Fig.5F). The high R² (coefficient of determination) value indicated a strong performance of the models in inferring CM maturation, highlighting the critical role of cellular communication in driving the CM maturation (Fig.5G). Moreover, the trained model highlighted not only the insulin pathway(*9*, *70*) but also *Ptn-Ncl*, the L-R pair for PTN pathway, as critical intercellular signaling mechanisms that contribute to CM maturation (Fig.5H, fig.S19).

Taken together, with our postnatal heart atlas, we uncovered the regulatory landscape underlying CM maturation, highlighting previously uncharacterized transcription factors and signaling pathways with potential roles in this process.

### Establishment of Probe-based Indel-detectable *in vivo* Perturb-seq for high-throughput functional screening at single-cell resolution

Next, we sought to establish a high-throughput *in vivo* system to assess the role of uncharacterized regulators in CM maturation, as determined by our integrative analysis. Compared to conventional knockout mouse models, while widely used but inherently low throughput, in vivo Perturb-seq platforms provide a high-throughput alternative to examine gene’s function within the complex physiological dynamics of the *in vivo* environment(*29*, *71*, *72*). Although published *in vivo* Perturb-seq platforms have been optimized to read out perturbed transcriptomes at the single-cell level, their reliance on droplet-based single-cell library preparation makes them challenging to apply directly to cardiomyocytes, which, due to their large size, is not suitable for droplet formation. To overcome this limitation, we adopted the conceptual framework of *in vivo* Perturb-seq and developed a modified version that utilizes single nucleus as input. This adaptation enables simultaneous gene perturbation and functional assessment of targeted CMs within their native microenvironment (Fig. 6A).

**Figure 6.**
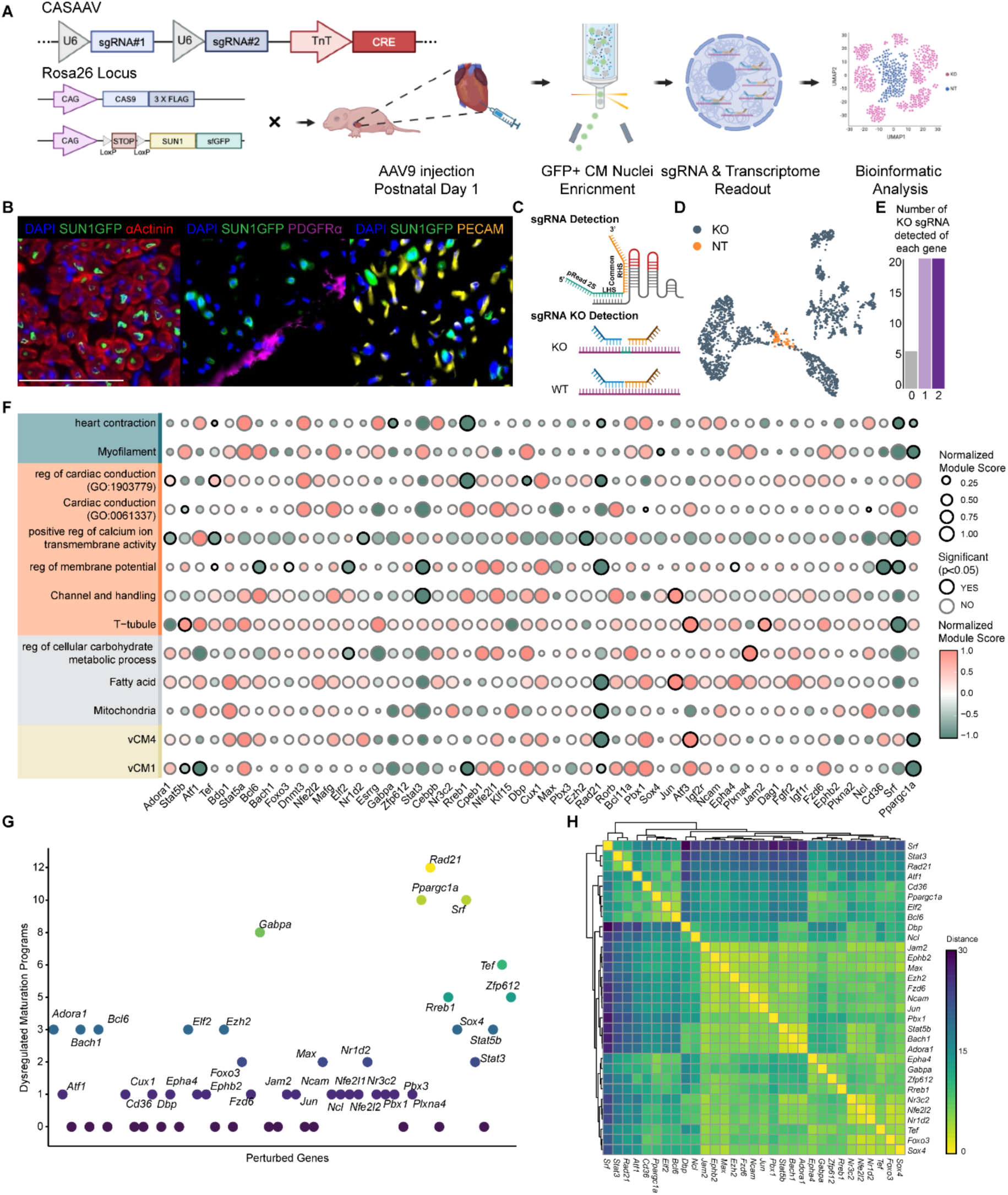
Establishment of Probe-based Indel-detectable *in vivo* Perturb-seq for high-throughput functional screening at single-cell resolution. A) Schematic showing the workflow of *in vivo* Probe-based Indel-detectable Perturb-seq. B) Fluorescence images showing the co-staining of SUN1-GFP (green) with other cell type makers, αActinin (left panel, red), PDFGRα (Middle panel, purple), and PECAM1 (right panel, yellow). Scale bar, 75μm. C) Schematic showing the design strategy for sgRNA detection probe (Top) and sgRNA KO detection probe (Bottom). D) UMAP of perturbed cells (navel blue) and nontargeting cells (orange). E) Bar plot showing the distribution of the number of sgRNAs for each gene with successful knockout detection using sgRNA KO detection probe. F) Dot plot showing each perturbation’s effect on expression of genes in maturation-related program (Green: Myofibril, Orange: Electrophysiology, Grey: Metabolism, Yellow: WGCNA module). Wilcoxon test was performed between perturbed cells and non-targeting cells. Gene sets showing statistically significant perturbation between perturbed cells and non-targeting cells were labeled with a black border. G) Scatter plot summarizing the number of significantly dysregulated maturation programs by each perturbed gene. H) Heatmap showing the sample distance between each sgRNA’s impact on the maturation programs. Color scheme represents the sample distance from high (yellow) to low (dark blue).

For efficient and specific delivery of sgRNAs to postnatal mice, we utilized a AAV9 viral vector to carry a cTnT-promoter-driven Cre recombinase and two U6-promoter-driven sgRNA cassettes(*73*). This design ensures precise co-delivery of Cre recombinase and the two sgRNAs, which target the same individual candidate CM maturation regulators, to CMs (Fig.6A). Additionally, neonatal mice receiving the AAV9 viruses harbored two transgenes: one constitutively expressing Cas9 to support sgRNA-guided gene knockout and another containing a floxed-stop cassette followed by nuclear membrane-anchored SUN1-sfGFP fusion protein. Upon Cre-mediated excision of the floxed STOP cassette, this system specifically labels sgRNA-expressing CMs through SUN1-sfGFP localization on the nuclear membrane (fig.S20A). The effectiveness of *in vivo* gene editing was confirmed using T7E1 assay (fig.S20B), and CM-specific labeling was validated by co-staining SUN1-sfGFP with CM and non-CM markers (Fig.6B).

To simultaneously detect sgRNA expression and comprehensively profile cardiac transcriptome at the single nucleus level, we adapted the Single Cell Gene Expression Flex assay from 10x Genomics. Compared to standard snRNA-seq, this fixation-based method mitigates dissociation-induced transcriptional stress responses and preserves RNA integrity by stabilizing transcripts within intact nuclei before processing. Furthermore, the probe-based chemistry of fixed RNA profiling supports the design of custom probes for each sgRNA, facilitating direct detection of the sgRNA expression (Fig.6C). With its high sensitivity, the Flex assay significantly enhances sgRNA capture efficiency compared to the unfixed RNA profiling methods (fig.S20C). This advantage is even more pronounced in intact nuclei, where RNA abundance is substantially lower than in whole cells. Additionally, the sequence specificity of probe hybridization allows for accurate determination of perturbation states in individual targeted cells. The perturbation detection probes are designed to capture wild-type transcripts at sgRNA target sites. These probes hybridize specifically to intact RNA molecules but fail to bind when Cas9-induced indels disrupt the target sequence (Fig. 6C). Consequently, cells with successful gene editing exhibit lower transcript counts compared to the non-targeting group, indicating target gene perturbation. Together, these features establish Probe-based Indel-detectable *in vivo* Perturb-seq (PIP-seq) as a powerful platform for high-throughput functional interrogation of candidate genes in a physiologically relevant context at single-cell resolution.

### Applying PIP-seq to identify novel regulators of cardiomyocyte maturation

We then applied PIP-seq to assess the feasibility of this approach in interrogating the putative regulators controlling CM maturation. The AAV9 library contains viral vectors encoding dual sgRNAs, designed to target 50 individual candidate regulators, including 37 intracellular and 13 intercellular ones, identified through our genomic analyses (fig.S21A, Supplementary Table.2). Previously validated non-targeting sgRNAs were included as the negative control, and sgRNAs targeting known regulators for CM maturation - *Srf*(*74*) and *Ppargc1a*(*23*) - are included as the positive control. To ensure a predominantly single AAV9 infection per GFP-positive CM, a multiplicity of infection (MOI) of 0.3 was selected based on a virus titration experiment (fig.S21B-D). Next, to determine the end point of our PIP-seq, we conducted permutation-based pairwise comparisons across maturation stages to assess the transcriptome dynamics of CMs (fig.S21E-F). Overall, CMs exhibited more significant transcriptome changes from P0 to P14 than from P14 to P21, indicating p14 as a critical time point when CMs attain a transcriptionally mature state, in line with our regulome analyses of the postnatal heart (Fig.3A-B). Therefore, neonatal hearts infected with AAV9 viruses at p0 were harvested at p14 for fixed RNA profiling to detect sgRNA identity and interrogate the transcriptomic alterations of FACS-sorted GFP+ nuclei at p14 in *in vivo* PIP-seq (fig.S22A-B).

A total of 5,706 nuclei, each with at least one assigned sgRNA, were recovered for downstream perturbation analysis (fig.S22C). Uniform Approximation and Projection (UAMP) plots demonstrated a clear separation between non-targeting CMs and perturbed nuclei (Fig.6D). This distinct clustering confirms effective gene perturbation and highlights the robustness of the *in vivo* PIP-seq platform in capturing cardiac transcriptomic alterations. To assess the knockout efficiency for each sgRNAs, we compared the unique molecular identification (UMI) read counts of the perturbation detection probes between perturbed and non-targeting control nuclei. Notably, 88% of the candidates have at least one sgRNA exhibiting reduced detection levels of the perturbation detection probes upon perturbation, highlighting the advantage of dual sgRNA design to achieve a high gene perturbation efficiency in the perturbed CMs (Fig. 6E).

To further determine the impact of each successful perturbation on the transcriptional program underlying CM maturation, we computed module scores to quantify the normalized expression levels of key gene sets associated with maturation. These gene sets comprised genes enriched in previously reported gene ontology (GO) associated with fundamental biological processes in CM maturation, genes demonstrating increased expression during CM maturation as identified in the snRNA-seq dataset, and gene modules derived from the WGCNA analysis that exhibit elevated expression throughout maturation. As expected, these gene sets showed increasing module scores along the postnatal CM maturation process (fig.S22D). Among the 50 candidate regulators analyzed, we identified 23 genes, including the established positive control *Srf* and *Ppargc1a,* that exerted a statistically significant effect on at least one of the transcriptional programs associated with CM maturation (Fig.6F-G).

Furthermore, we observed differential effects in the transcriptional response elicited by perturbation of these candidate maturation regulators, indicating that individual candidate regulators exert distinct modulatory effects on specific gene programs governing CM maturation (fig.S23). Targeting *Rad21,* a key component of the cohesin complex involved in chromatin organization and transcriptional regulation, resulted in a wide disruption of gene expression programs involved in myofibril organization, electrophysiological properties, and fatty acid metabolism, suggesting its broad role in coordinating multiple aspects of postnatal CM maturation (Fig.6F, fig.S23). Likewise, perturbing *Rreb1*, the Ras-responsive element-binding protein 1, led to marked disruptions in both myofibril organization and electrophysiological properties (Fig.6F, fig.S23). Subsequently, we investigated the relationship between each sgRNA knockout by assessing the overall similarity in the impact of each knockout on the maturation programs. Similarly to *Srf*, the master regulator of cardiomyocyte maturation, *Rad21* and *Stat3* exhibited a global impact on the maturation programs (Fig.6H). Notably, we also discovered that *Cd36* KO exhibited a remarkably similar impact on the maturation programs as *Ppargc1a*, precisely reflecting the involvement of CD36-PPARs Pathway in lipid metabolism (Fig.6H)(*75*). These findings underscore the distinct and collaborative efforts of these regulators in modulating the maturation of postnatal cardiomyocytes.

Having validated candidate maturation regulators at the transcriptional level using *in vivo* PIP-seq, we sought to determine whether the transcriptional alterations lead to phenotypic consequences. To functionally characterize the identified regulators, we performed targeted perturbation of *Rreb1*, whose role in CM maturation has not been discovered due to the embryonic lethality of its homozygous knockout(*76*). To bypass this limitation, we injected AAV9 viruses containing sgRNAs targeting *Rreb1* into p0 neonatal mice (Fig.7A). CMs successfully infected with the viruses were identified by GFP-labeled nuclear membranes (Fig.7B). At p14, CM size, a hallmark of CM maturation, was compared between the GFP + and GFP- CMs from the same heart to minimize the biological variability. Consistent with our PIP-seq findings, perturbation of *Rreb1* disrupted CM maturation, as indicated by reduced CM size compared to GFP⁻ counterparts (Fig. 7C-D). Notably, *Rreb1* perturbation specifically reduced the longitudinal size of CMs (Fig. 7E-F). Consistent with the cellular phenotype, *Rreb1* perturbation also leads to a significant reduction in heart size (Fig.7G). To further investigate the role of *Rreb1* in postnatal heart development, we performed echocardiography to assess heart function at p28, providing a higher dosage of the AAV- sgRreb1 and a longer duration for the phenotype to manifest (Fig.7H). *Rreb1* perturbation resulted in reduced heart function, characterized by the reduced fractional shortening and ejection fraction (Fig.7I-K).

**Figure 7.**
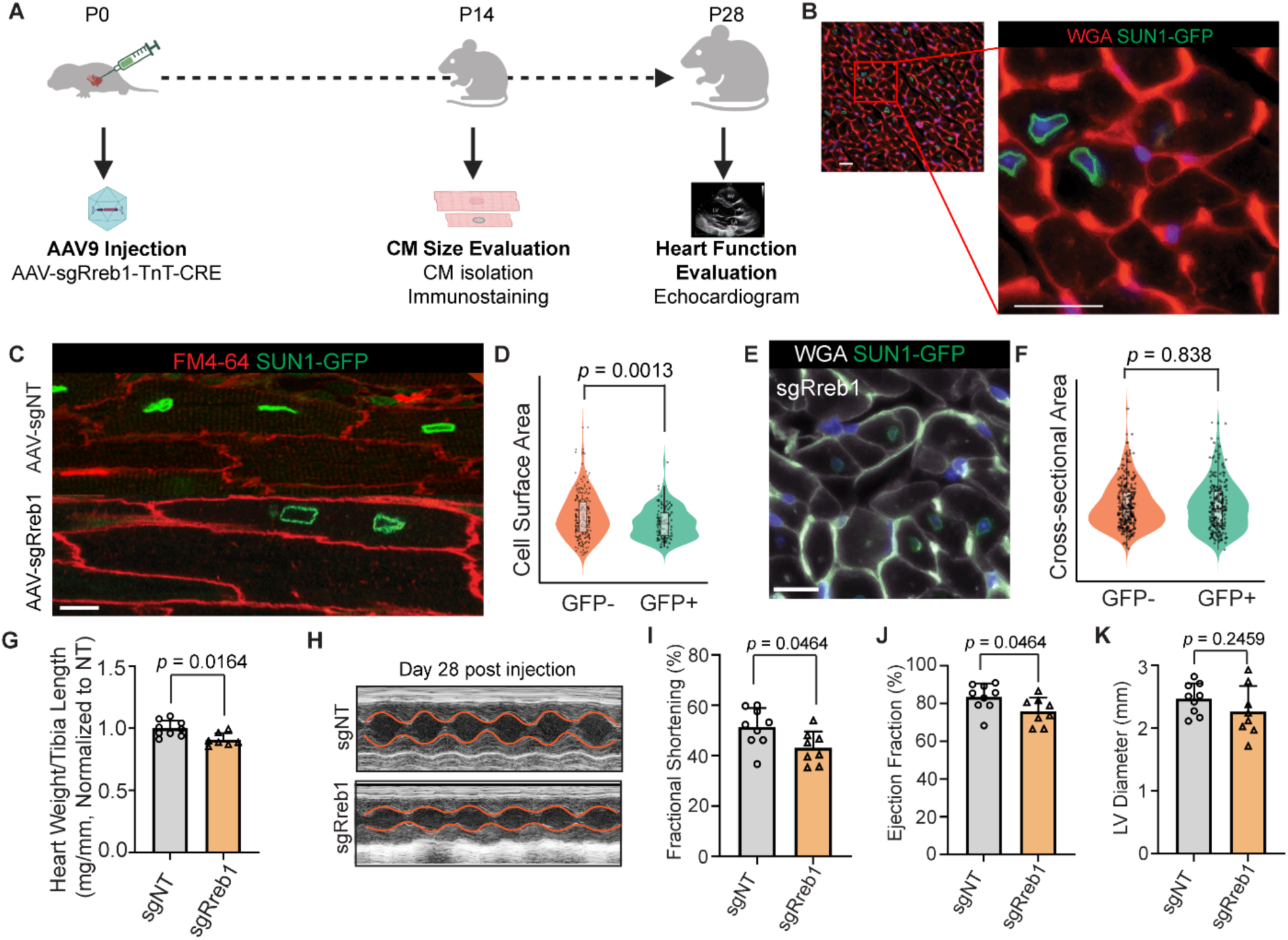
Targeted Knockout of *Rreb1* in the postnatal heart disrupted postnatal cardiomyocyte maturation. A) Schematic showing the workflow of validating the function of the PIP-seq candidate gene through a single AAV injection of the postnatal heart. B) Immunofluorescent image showing the infected cardiomyocyte is labeled with GFP at its nucleus membrane. Scale bar, 10 μm. C) Immunofluorescent image showing that the knockout of *Rreb1* leads to reduced cardiomyocyte cell size (longitudinal Area) compared to the GFP-negative cardiomyocyte. Scale bar, 10 μm. D) The quantitative analysis showing that the knockout of *Rreb1* leads to reduced cardiomyocyte cell size compared to the GFP-negative cardiomyocyte. E) Immunofluorescent image showing that the effect of knocking out *Rreb1* on the cardiomyocyte cell size (cross-section) compared to the GFP-negative cardiomyocyte. Scale bar, 10 μm. F) Quantitative analysis showing that the effect knocking out *Rreb1* on the cardiomyocyte cell size (cross-section) compared to the GFP-negative cardiomyocyte. G) Bar plot showing the effect of *Rreb1* knockout on Heart Weight versus Tibia Length. Independent samples t-test were performed to test the statistical significance of the Rreb1 knockout H) Echocardiogram showing the effect of *Rreb1* knockout on the postnatal heart function. I) Bar plot showing the effect of *Rreb1* knockout on the fractional shorting of the postnatal heart. Independent samples t-test were performed to test the statistical significance of the Rreb1 knockout. J) Bar plot showing the effect of *Rreb1* knockout on the ejection fraction of the postnatal heart. Independent samples t-test was performed to test the statistical significance of the Rreb1 knockout. K) Bar plot showing the effect of Rreb1’s knockout on the left ventricle diameter of the postnatal heart. Independent samples t-test was performed to test the statistical significance of the *Rreb1* knockout.

Overall, our *in vivo* PIP-seq platform demonstrate its capability to identify functional regulators for the biological process of interest in a high-throughput manner, all within an *in vivo* physiology context. Applying this platform to the postnatal CMs, we identified 21 novel regulators involved in regulating distinct aspects of postnatal CM maturation transcriptional programs.

## Discussion

In this study, we generated a high-resolution single-cell atlas of the postnatal murine heart, capturing dynamic changes in cell composition and gene expression across critical postnatal developmental stages. By integrating single-nucleus RNA sequencing with Xenium spatial transcriptomics, we delineated the temporal dynamics of cardiac cell types and their spatially defined cellular neighborhoods, revealing their associations with distinct anatomical structures. This approach enabled the identification of intrinsic and extrinsic regulatory mechanisms governing postnatal heart development, including novel transcriptional regulons and ligand-receptor interactions. Notably, we uncovered a key interaction between cap.ECs and postnatal CMs, which appears to drive CM maturation by promoting cell cycle exit. Temporal analysis of CM transcriptome dynamics further constructed a comprehensive regulatory network, identifying both known and previously uncharacterized regulators of CM maturation. Additionally, our analysis of postnatal CM interactome elucidates the stage-specific contributions of various signaling pathways to postnatal cardiac development.

To functionally interrogate these newly identified regulators within their physiological context, we developed PIP-seq, an optimized *in vivo* Perturb-seq platform. By leveraging probe-based fixed RNA profiling, PIP-seq enables direct detection of sgRNA expression and perturbation effects at single-cell resolution. Unlike conventional *in vivo* Perturb-seq, which relies on single-cell dissociation, PIP-seq employs single-nucleus profiling, making it particularly suited for cell types that are difficult to dissociate or sort, such as CMs and hepatocytes. Additionally, PIP-seq’s probe-based chemistry facilitates the precise determination of perturbation states, allowing for the exclusion of non-perturbed cells and ineffective sgRNAs. Its mosaic perturbation strategy further enables gene function studies in specific biological processes while mitigating secondary effects associated with whole-organ or systemic perturbations. Using PIP-seq, we identified 23 regulators of CM maturation, including *Rreb1*, whose role in postnatal CM maturation had remained unexplored due to the embryonic lethality of its homozygous knockout model. Functional validation of *Rreb1* perturbation confirmed its role in CM maturation and cardiac function, further demonstrating the physiological relevance of our approach.

Overall, our integrative approach provides an unprecedented framework for studying postnatal heart development at high spatial and temporal resolution. By bridging large-scale transcriptomic mapping with high-throughput *in vivo* functional screening, our study not only deepens our understanding of cardiac maturation but also establishes PIP-seq as a powerful tool for investigating gene function in complex biological systems.

## Supporting information

Suppementary datas

## Acknowledgements

We thank the UNC Vector Core Facility for packaging the AAV virus. We thank the UNC High-Throughput Sequencing Facility for all sequencing work and assistance on bioinformatics analysis. We thank the UNC Flow Cytometry Core Facility for nuclei sorting. Figure 1A, 1F, 2D, 3H, 5A, 6C, and 7A are created in BioRender. During the preparation of this work, the authors used generative AI in order to refine grammar and language. The authors reviewed and edited the content as needed and take full responsibility for the content of the publication.

## Author Contributions

Conceptualization: H.W., L.Q., and J.L. Methodology: H.W., Y.D., and Y.S. Investigation: H.W., Y.D., Y.S., M.C., N.Y., S.R., X.L., G.F., and Y.Q. Visualization: H.W., Y.D., Y.S., and M.C., Supervision: J.L. and L.Q. Writing—original draft: H.W., Y.D., L.Q., J.L. Writing—review and editing: H.W., Y.D., Y.S., M.C., N.Y., S.N., X.L., G.F., Y.Q., L.Q., and J.L.

## Data and materials Availability

Sequencing data will be deposited at GEO. Our codebase for the figure generation and PIP-seq pipeline and perturbation analysis will be available on GitHub.

## References and Notes

1. A. Rao, D. Barkley, G. S. França, I. Yanai, Exploring tissue architecture using spatial transcriptomics. Nature 596, 211–220 (2021).

2. B. L. Walker, Z. Cang, H. Ren, E. Bourgain-Chang, Q. Nie, Deciphering tissue structure and function using spatial transcriptomics. Commun Biol 5, 1–10 (2022).

3. J. L. Close, B. R. Long, H. Zeng, Spatially resolved transcriptomics in neuroscience. Nat Methods 18, 23–25 (2021).

4. X. Zhuang, Spatially resolved single-cell genomics and transcriptomics by imaging. Nat Methods 18, 18–22 (2021).

5. D. Bressan, G. Battistoni, G. J. Hannon, The dawn of spatial omics. Science 381, eabq4964 (2023).

6. K. Kanemaru, J. Cranley, D. Muraro, A. M. A. Miranda, S. Y. Ho, A. Wilbrey-Clark, J. Patrick Pett, K. Polanski, L. Richardson, M. Litvinukova, N. Kumasaka, Y. Qin, Z. Jablonska, C. I. Semprich, L. Mach, M. Dabrowska, N. Richoz, L. Bolt, L. Mamanova, R. Kapuge, S. N. Barnett, S. Perera, C. Talavera-López, I. Mulas, K. T. Mahbubani, L. Tuck, L. Wang, M. M. Huang, M. Prete, S. Pritchard, J. Dark, K. Saeb-Parsy, M. Patel, M. R. Clatworthy, N. Hübner, R. A. Chowdhury, M. Noseda, S. A. Teichmann, Spatially resolved multiomics of human cardiac niches. Nature 619, 801–810 (2023).

7. E. N. Farah, R. K. Hu, C. Kern, Q. Zhang, T.-Y. Lu, Q. Ma, S. Tran, B. Zhang, D. Carlin, A. Monell, A. P. Blair, Z. Wang, J. Eschbach, B. Li, E. Destici, B. Ren, S. M. Evans, S. Chen, Q. Zhu, N. C. Chi, Spatially organized cellular communities form the developing human heart. Nature 627, 854–864 (2024).

8. J. Cranley, S. Bayraktar, K. Kanemaru, V. Knight-Schrijver, J. P. Pett, H. Davaapil, L. Gambardella, S. Sinha, S. Teichmann, A spatially-resolved multiomic cell atlas reveals gene regulatory networks underlying cell specification in the developing human heart. European Heart Journal 44 (2023).

9. T. A. Garbutt, Z. Wang, H. Wang, H. Ma, H. Ruan, Y. Dong, Y. Xie, L. Tan, R. Phookan, J. A. Stouffer, V. Vedantham, Y. Yang, L. Qian, J. Liu, Epigenetic Regulation of Cardiomyocyte Maturation by Arginine Methyltransferase CARM1. Circulation 149, 1501–1515 (2024).

10. N. Velayutham, E. J. Agnew, K. E. Yutzey, Postnatal cardiac development and regenerative potential in large mammals. Pediatr Cardiol 40, 1345–1358 (2019).

11. V. Talman, J. Teppo, P. Pöhö, P. Movahedi, A. Vaikkinen, S. T. Karhu, K. Trošt, T. Suvitaival, J. Heikkonen, T. Pahikkala, T. Kotiaho, R. Kostiainen, M. Varjosalo, H. Ruskoaho, Molecular Atlas of Postnatal Mouse Heart Development. J Am Heart Assoc 7, e010378 (2018).

12. T. Sada, W. Kimura, Transition from fetal to postnatal state in the heart: Crosstalk between metabolism and regeneration. Development, Growth & Differentiation 66, 438–451 (2024).

13. N. J. VanDusen, J. Y. Lee, W. Gu, C. E. Butler, I. Sethi, Y. Zheng, J. S. King, P. Zhou, S. Suo, Y. Guo, Q. Ma, G.-C. Yuan, W. T. Pu, Massively parallel in vivo CRISPR screening identifies RNF20/40 as epigenetic regulators of cardiomyocyte maturation. Nat Commun 12, 4442 (2021).

14. Y. Guo, W. T. Pu, Cardiomyocyte Maturation. Circulation Research, 1086–1106 (2020).

15. S. L. Padula, N. Velayutham, K. E. Yutzey, Transcriptional Regulation of Postnatal Cardiomyocyte Maturation and Regeneration. Int J Mol Sci 22, 3288 (2021).

16. T. Sakamoto, D. P. Kelly, Cardiac maturation. Journal of Molecular and Cellular Cardiology 187, 38–50 (2024).

17. S. Salameh, V. Ogueri, N. G. Posnack, Adapting to a new environment: postnatal maturation of the human cardiomyocyte. J Physiol 601, 2593–2619 (2023).

18. P. Hu, J. Liu, J. Zhao, B. J. Wilkins, K. Lupino, H. Wu, L. Pei, Single-nucleus transcriptomic survey of cell diversity and functional maturation in postnatal mammalian hearts. Genes Dev 32, 1344–1357 (2018).

19. Z. Li, F. Yao, P. Yu, D. Li, M. Zhang, L. Mao, X. Shen, Z. Ren, L. Wang, B. Zhou, Postnatal state transition of cardiomyocyte as a primary step in heart maturation. Protein Cell 13, 842– 862 (2022).

20. Y. Wang, F. Yao, L. Wang, Z. Li, Z. Ren, D. Li, M. Zhang, L. Han, S. Wang, B. Zhou, L. Wang, Single-cell analysis of murine fibroblasts identifies neonatal to adult switching that regulates cardiomyocyte maturation. Nature Communications 11, 2585 (2020).

21. W. Feng, A. Bais, H. He, C. Rios, S. Jiang, J. Xu, C. Chang, D. Kostka, G. Li, Single-cell transcriptomic analysis identifies murine heart molecular features at embryonic and neonatal stages. Nature Communications 2022 13:*1* 13, 1–19 (2022).

22. D. A. Skelly, G. T. Squiers, M. A. McLellan, M. T. Bolisetty, P. Robson, N. A. Rosenthal, A. R. Pinto, Single-Cell Transcriptional Profiling Reveals Cellular Diversity and Intercommunication in the Mouse Heart. Cell Reports 22, 600–610 (2018).

23. S. A. Murphy, M. Miyamoto, A. Kervadec, S. Kannan, E. Tampakakis, S. Kambhampati, B. L. Lin, S. Paek, P. Andersen, D. I. Lee, R. Zhu, S. S. An, D. A. Kass, H. Uosaki, A. R. Colas, C. Kwon, PGC1/PPAR drive cardiomyocyte maturation at single cell level via YAP1 and SF3B2. Nature Communications 2021 12:*1* 12, 1–12 (2021).

24. S. Kannan, M. Miyamoto, R. Zhu, M. Lynott, J. Guo, E. Z. Chen, A. R. Colas, B. L. Lin, C. Kwon, Trajectory reconstruction identifies dysregulation of perinatal maturation programs in pluripotent stem cell-derived cardiomyocytes. Cell Reports 42, 112330 (2023).

25. I. R. Brooks, C. M. Garrone, C. Kerins, C. S. Kiar, S. Syntaka, J. Z. Xu, F. M. Spagnoli, F. M. Watt, Functional genomics and the future of iPSCs in disease modeling. Stem Cell Reports 17, 1033–1047 (2022).

26. A. Padmanabhan, T. Y. de Soysa, A. Pelonero, V. Sapp, P. P. Shah, Q. Wang, L. Li, C. Y. Lee, N. Sadagopan, T. Nishino, L. Ye, R. Yang, A. Karnay, A. Poleshko, N. Bolar, R. Linares-Saldana, S. S. Ranade, M. Alexanian, S. U. Morton, M. Jain, S. M. Haldar, D. Srivastava, R. Jain, A genome-wide CRISPR screen identifies BRD4 as a regulator of cardiomyocyte differentiation. Nat Cardiovasc Res 3, 317–331 (2024).

27. P. Datlinger, A. F. Rendeiro, C. Schmidl, T. Krausgruber, P. Traxler, J. Klughammer, L. C. Schuster, A. Kuchler, D. Alpar, C. Bock, Pooled CRISPR screening with single-cell transcriptome readout. Nat Methods 14, 297–301 (2017).

28. J. M. Replogle, R. A. Saunders, A. N. Pogson, J. A. Hussmann, A. Lenail, A. Guna, L. Mascibroda, E. J. Wagner, K. Adelman, G. Lithwick-Yanai, N. Iremadze, F. Oberstrass, D. Lipson, J. L. Bonnar, M. Jost, T. M. Norman, J. S. Weissman, Mapping information-rich genotype-phenotype landscapes with genome-scale Perturb-seq. Cell 185, 2559–2575.e28 (2022).

29. X. Jin, S. K. Simmons, A. Guo, A. S. Shetty, M. Ko, L. Nguyen, V. Jokhi, E. Robinson, P. Oyler, N. Curry, G. Deangeli, S. Lodato, J. Z. Levin, A. Regev, F. Zhang, P. Arlotta, In vivo Perturb-Seq reveals neuronal and glial abnormalities associated with autism risk genes. Science 370, eaaz6063 (2020).

30. N. J. VanDusen, J. Y. Lee, W. Gu, C. E. Butler, I. Sethi, Y. Zheng, J. S. King, P. Zhou, S. Suo, Y. Guo, Q. Ma, G. C. Yuan, W. W. Pu, Massively parallel in vivo CRISPR screening identifies RNF20/40 as epigenetic regulators of cardiomyocyte maturation. Nature communications 12 (2021).

31. L. Machado, F. Relaix, P. Mourikis, Stress relief: emerging methods to mitigate dissociation-induced artefacts. Trends in Cell Biology 31, 888–897 (2021).

32. M. R. Vahid, E. L. Brown, C. B. Steen, W. Zhang, H. S. Jeon, M. Kang, A. J. Gentles, A. M. Newman, High-resolution alignment of single-cell and spatial transcriptomes with CytoSPACE. Nat Biotechnol 41, 1543–1548 (2023).

33. Y.-C. Chen, A. Suresh, C. Underbayev, C. Sun, K. Singh, F. Seifuddin, A. Wiestner, M. Pirooznia, IKAP-Identifying K mAjor cell Population groups in single-cell RNA-sequencing analysis. Gigascience 8, giz121 (2019).

34. N. P. Murphy, E. R. Lubbers, P. J. Mohler, Advancing our understanding of AnkRD1 in cardiac development and disease. Cardiovascular Research 116, 1402–1404 (2020).

35. L. Zhong, M. Chiusa, A. G. Cadar, A. Lin, S. Samaras, J. M. Davidson, C. C. Lim, Targeted inhibition of ANKRD1 disrupts sarcomeric ERK-GATA4 signal transduction and abrogates phenylephrine-induced cardiomyocyte hypertrophy. Cardiovasc Res 106, 261–271 (2015).

36. S. Boskovic, R. Marín Juez, N. Stamenkovic, D. Radojkovic, D. Y. Stainier, S. Kojic, The stress responsive gene ankrd1a is dynamically regulated during skeletal muscle development and upregulated following cardiac injury in border zone cardiomyocytes in adult zebrafish. Gene 792, 145725 (2021).

37. C. Kuppe, R. O. Ramirez Flores, Z. Li, S. Hayat, R. T. Levinson, X. Liao, M. T. Hannani, J. Tanevski, F. Wünnemann, J. S. Nagai, M. Halder, D. Schumacher, S. Menzel, G. Schäfer, K. Hoeft, M. Cheng, S. Ziegler, X. Zhang, F. Peisker, N. Kaesler, T. Saritas, Y. Xu, A. Kassner, J. Gummert, M. Morshuis, J. Amrute, R. J. A. Veltrop, P. Boor, K. Klingel, L. W. Van Laake, A. Vink, R. M. Hoogenboezem, E. M. J. Bindels, L. Schurgers, S. Sattler, D. Schapiro, R. K. Schneider, K. Lavine, H. Milting, I. G. Costa, J. Saez-Rodriguez, R. Kramann, Spatial multi-omic map of human myocardial infarction. Nature 608, 766–777 (2022).

38. T. Hai, C. D. Wolfgang, D. K. Marsee, A. E. Allen, U. Sivaprasad, ATF3 and Stress Responses. Gene Expr 7, 321–335 (2018).

39. N. Yucel, J. Axsom, Y. Yang, L. Li, J. H. Rhoades, Z. Arany, Cardiac endothelial cells maintain open chromatin and expression of cardiomyocyte myofibrillar genes. eLife 9, 1–19 (2020).

40. R. G. Li, X. Li, Y. Morikawa, F. J. Grisanti-Canozo, F. Meng, C.-R. Tsai, Y. Zhao, L. Liu, J. Kim, B. Xie, E. Klysik, S. Liu, M. A. H. Samee, J. F. Martin, YAP induces a neonatal-like pro-renewal niche in the adult heart. Nat Cardiovasc Res 3, 283–300 (2024).

41. Z. Cang, Y. Zhao, A. A. Almet, A. Stabell, R. Ramos, M. V. Plikus, S. X. Atwood, Q. Nie, Screening cell-cell communication in spatial transcriptomics via collective optimal transport. Nat Methods 20, 218–228 (2023).

42. M. Ieda, H. Kanazawa, K. Kimura, F. Hattori, Y. Ieda, M. Taniguchi, J.-K. Lee, K. Matsumura, Y. Tomita, S. Miyoshi, K. Shimoda, S. Makino, M. Sano, I. Kodama, S. Ogawa, K. Fukuda, Sema3a maintains normal heart rhythm through sympathetic innervation patterning. Nat Med 13, 604–612 (2007).

43. O. Salvucci, G. Tosato, Essential Roles of EphB Receptors and EphrinB Ligands in Endothelial Cell Function and Angiogenesis. Adv Cancer Res 114, 21–57 (2012).

44. T. Ishida, R. K. Kundu, E. Yang, K. Hirata, Y.-D. Ho, T. Quertermous, Targeted disruption of endothelial cell-selective adhesion molecule inhibits angiogenic processes in vitro and in vivo. J Biol Chem 278, 34598–34604 (2003).

45. M. Segarra, H. Ohnuki, D. Maric, O. Salvucci, X. Hou, A. Kumar, X. Li, G. Tosato, Semaphorin 6A regulates angiogenesis by modulating VEGF signaling. Blood 120, 4104– 4115 (2012).

46. M. Hellström, L.-K. Phng, J. J. Hofmann, E. Wallgard, L. Coultas, P. Lindblom, J. Alva, A.-K. Nilsson, L. Karlsson, N. Gaiano, K. Yoon, J. Rossant, M. L. Iruela-Arispe, M. Kalén, H. Gerhardt, C. Betsholtz, Dll4 signalling through Notch1 regulates formation of tip cells during angiogenesis. Nature 445, 776–780 (2007).

47. Z.-J. Liu, T. Shirakawa, Y. Li, A. Soma, M. Oka, G. P. Dotto, R. M. Fairman, O. C. Velazquez, M. Herlyn, Regulation of Notch1 and Dll4 by Vascular Endothelial Growth Factor in Arterial Endothelial Cells: Implications for Modulating Arteriogenesis and Angiogenesis. Mol Cell Biol 23, 14–25 (2003).

48. P. Lu, Y. Wang, Y. Liu, Y. Wang, B. Wu, D. Zheng, R. P. Harvey, B. Zhou, Perinatal angiogenesis from pre-existing coronary vessels via DLL4-NOTCH1 signalling. Nat Cell Biol 23, 967–977 (2021).

49. M. Shibuya, Vascular Endothelial Growth Factor (VEGF) and Its Receptor (VEGFR) Signaling in Angiogenesis. Genes Cancer 2, 1097–1105 (2011).

50. J. Matsui, T. Wakabayashi, M. Asada, K. Yoshimatsu, M. Okada, Stem Cell Factor/c-kit Signaling Promotes the Survival, Migration, and Capillary Tube Formation of Human Umbilical Vein Endothelial Cells *. Journal of Biological Chemistry 279, 18600–18607 (2004).

51. S. Aibar, C. B. González-Blas, T. Moerman, V. A. Huynh-Thu, H. Imrichova, G. Hulselmans, F. Rambow, J. C. Marine, P. Geurts, J. Aerts, J. Van Den Oord, Z. K. Atak, J. Wouters, S. Aerts, SCENIC: single-cell regulatory network inference and clustering. Nature methods 14, 1083–1086 (2017).

52. C. Schmal, J. Myung, H. Herzel, G. Bordyugov, Moran’s I quantifies spatio-temporal pattern formation in neural imaging data. Bioinformatics 33, 3072–3079 (2017).

53. S. Seo, T. Kume, Forkhead transcription factors, Foxc1 and Foxc2, are required for the morphogenesis of the cardiac outflow tract. Dev Biol 296, 421–436 (2006).

54. E. Lambers, B. Arnone, A. Fatima, G. Qin, J. A. Wasserstrom, T. Kume, Foxc1 Regulates Early Cardiomyogenesis and Functional Properties of Embryonic Stem Cell Derived Cardiomyocytes. STEM CELLS 34, 1487–1500 (2016).

55. L. He, Q. Zhang, D. Jiang, Y. Zhang, Y. Wei, Y. Yang, N. Li, S. Wang, Y. Yue, Q. Zhao, Zebrafish Foxc1a controls ventricular chamber maturation by directly regulating *wwtr1* and *nkx2.5* expression. Journal of Genetics and Genomics 49, 559–568 (2022).

56. J. W. Squair, M. A. Skinnider, M. Gautier, L. J. Foster, G. Courtine, Prioritization of cell types responsive to biological perturbations in single-cell data with Augur. Nat Protoc 16, 3836– 3873 (2021).

57. P.-H. Sun, L. Ye, M. D. Mason, W. G. Jiang, Protein Tyrosine Phosphatase µ (PTP µ or PTPRM), a Negative Regulator of Proliferation and Invasion of Breast Cancer Cells, Is Associated with Disease Prognosis. PLOS ONE 7, e50183 (2012).

58. A. J. Hale, E. ter Steege, J. den Hertog, Recent advances in understanding the role of protein-tyrosine phosphatases in development and disease. Developmental Biology 428, 283–292 (2017).

59. A. I. Mahmoud, F. Kocabas, S. A. Muralidhar, W. Kimura, A. S. Koura, S. Thet, E. R. Porrello, H. A. Sadek, Meis1 regulates postnatal cardiomyocyte cell cycle arrest. Nature 497, 249–253 (2013).

60. S. Morabito, F. Reese, N. Rahimzadeh, E. Miyoshi, V. Swarup, hdWGCNA identifies co-expression networks in high-dimensional transcriptomics data. Cell Reports Methods 3, 100498 (2023).

61. N. N. Chattergoon, G. D. Giraud, S. Louey, P. Stork, A. L. Fowden, K. L. Thornburg, Thyroid hormone drives fetal cardiomyocyte maturation. FASEB journal : official publication of the Federation of American Societies for Experimental Biology 26, 397–408 (2012).

62. M. Li, S. E. Iismaa, N. Naqvi, A. Nicks, A. Husain, R. M. Graham, Thyroid hormone action in postnatal heart development. Stem Cell Res 13, 582–591 (2014).

63. G. C. Rowe, A. Jiang, Z. Arany, PGC-1 coactivators in cardiac development and disease. Circ Res 107, 825–838 (2010).

64. T. Sakamoto, T. R. Matsuura, S. Wan, D. M. Ryba, J. U. Kim, K. J. Won, L. Lai, C. Petucci, N. Petrenko, K. Musunuru, R. B. Vega, D. P. Kelly, A Critical Role for Estrogen-Related Receptor Signaling in Cardiac Maturation. Circ Res 126, 1685–1702 (2020).

65. S. Díaz Del Moral, M. Benaouicha, R. Muñoz-Chápuli, R. Carmona, The Insulin-like Growth Factor Signalling Pathway in Cardiac Development and Regeneration. International journal of molecular sciences 23 (2021).

66. J. Grego-Bessa, P. Gómez-Apiñaniz, B. Prados, M. J. Gómez, D. MacGrogan, J. L. de la Pompa, Nrg1 Regulates Cardiomyocyte Migration and Cell Cycle in Ventricular Development. Circ Res 133, 927–943 (2023).

67. O. Odiete, M. F. Hill, D. B. Sawyer, Neuregulin in Cardiovascular Development and Disease. Circ Res 111, 1376–1385 (2012).

68. Y. Y. Zhao, D. R. Sawyer, R. R. Baliga, D. J. Opel, X. Han, M. A. Marchionni, R. A. Kelly, Neuregulins promote survival and growth of cardiac myocytes. Persistence of ErbB2 and ErbB4 expression in neonatal and adult ventricular myocytes. J Biol Chem **273**, 10261–10269 (1998).

69. H. Ma, C. Yin, Y. Zhang, L. Qian, J. Liu, ErbB2 is required for cardiomyocyte proliferation in murine neonatal hearts. Gene 592, 325–330 (2016).

70. D. D. Belke, S. Betuing, M. J. Tuttle, C. Graveleau, M. E. Young, M. Pham, D. Zhang, R. C. Cooksey, D. A. McClain, S. E. Litwin, H. Taegtmeyer, D. Severson, C. R. Kahn, E. D. Abel, Insulin signaling coordinately regulates cardiac size, metabolism, and contractile protein isoform expression. J Clin Invest 109, 629–639 (2002).

71. A. J. Santinha, E. Klingler, M. Kuhn, R. Farouni, S. Lagler, G. Kalamakis, U. Lischetti, D. Jabaudon, R. J. Platt, Transcriptional linkage analysis with in vivo AAV-Perturb-seq. Nature 622, 367–375 (2023).

72. X. Zheng, B. Wu, Y. Liu, S. K. Simmons, K. Kim, G. S. Clarke, A. Ashiq, J. Park, J. Li, Z. Wang, L. Tong, Q. Wang, K. T. Rajamani, R. Muñoz-Castañeda, S. Mu, T. Qi, Y. Zhang, Z. C. Ngiam, N. Ohte, C. Hanashima, Z. Wu, X. Xu, J. Z. Levin, X. Jin, Massively parallel in vivo Perturb-seq reveals cell-type-specific transcriptional networks in cortical development. Cell, doi: 10.1016/J.CELL.2024.04.050 (2024).

73. N. J. VanDusen, Y. Guo, W. Gu, W. T. Pu, CASAAV: a CRISPR based platform for rapid dissection of gene function in vivo. Curr Protoc Mol Biol 120, 31.11.1–31.11.14 (2017).

74. Y. Guo, Y. Cao, B. D. Jardin, I. Sethi, Q. Ma, B. Moghadaszadeh, E. C. Troiano, N. Mazumdar, M. A. Trembley, E. M. Small, G.-C. Yuan, A. H. Beggs, W. T. Pu, Sarcomeres regulate murine cardiomyocyte maturation through MRTF-SRF signaling. Proc Natl Acad Sci U S A 118, e2008861118 (2021).

75. L. Maréchal, M. Laviolette, A. Rodrigue-Way, B. Sow, M. Brochu, V. Caron, A. Tremblay, The CD36-PPARγ Pathway in Metabolic Disorders. Int J Mol Sci 19, 1529 (2018).

76. O. A. Kent, M. Saha, E. Coyaud, H. E. Burston, N. Law, K. Dadson, S. Chen, E. M. Laurent, J. St-Germain, R. X. Sun, Y. Matsumoto, J. Cowen, A. Montgomery-Song, K. R. Brown, C. Ishak, J. L. Rose, D. D. De Carvalho, H. H. He, B. Raught, F. Billia, P. Kannu, R. Rottapel, Haploinsufficiency of RREB1 causes a Noonan-like RASopathy via epigenetic reprogramming of RAS-MAPK pathway genes. Nat Commun 11, 4673 (2020).

77. P.-L. Germain, A. Lun, C. Garcia Meixide, W. Macnair, M. D. Robinson, Doublet identification in single-cell sequencing data using scDblFinder. F1000Res **10**, 979 (2022).

78. T. Moerman, S. Aibar Santos, C. Bravo González-Blas, J. Simm, Y. Moreau, J. Aerts, S. Aerts, GRNBoost2 and Arboreto: efficient and scalable inference of gene regulatory networks. Bioinformatics 35, 2159–2161 (2019).

79. G. Palla, H. Spitzer, M. Klein, D. Fischer, A. C. Schaar, L. B. Kuemmerle, S. Rybakov, I. L. Ibarra, O. Holmberg, I. Virshup, M. Lotfollahi, S. Richter, F. J. Theis, Squidpy: a scalable framework for spatial omics analysis. Nat Methods 19, 171–178 (2022).

80. T. Li, D. Horsfall, D. Basurto-Lozada, K. Roberts, M. Prete, J. E. G. Lawrence, P. He, E. Tuck, J. Moore, A. K. Yoldas, K. Babalola, M. Hartley, S. Ghazanfar, S. A. Teichmann, M. Haniffa, O. A. Bayraktar, WebAtlas pipeline for integrated single-cell and spatial transcriptomic data. Nature methods, doi: 10.1038/S41592-024-02371-X (2024).

81. M. Ackers-Johnson, P. Y. Li, A. P. Holmes, S.-M. O’Brien, D. Pavlovic, R. S. Foo, A Simplified, Langendorff-Free Method for Concomitant Isolation of Viable Cardiac Myocytes and Nonmyocytes From the Adult Mouse Heart. Circ Res 119, 909–920 (2016).

82. Y. Guo, N. J. VanDusen, L. Zhang, W. Gu, I. Sethi, S. Guatimosim, Q. Ma, B. D. Jardin, Y. Ai, D. Zhang, B. Chen, A. Guo, G.-C. Yuan, L.-S. Song, W. T. Pu, Analysis of Cardiac Myocyte Maturation Using CASAAV, a Platform for Rapid Dissection of Cardiac Myocyte Gene Function In Vivo. Circulation Research 120, 1874–1888 (2017).

