## Supplementary material for "Charting Postnatal Heart Development Using *In Vivo* Single-Cell Functional Genomics": Suppementary datas

### Materials and Methods

#### Animals

Male and female wildtype, B6;129(Cg)-Gt(ROSA)26Sor$^{\text{tm1.1(CAG-cas9*,-EGFP)Fezh}}$/J in a C57BL/6J background, and B6;129-Gt(ROSA)26Sor$^{\text{tm5(CAG-Sun1/sfGFP)Nat}}$/J in C57BL/6 background were obtained from jackson laboratory and used in the current study. Animal care and experiments were carried out in accordance with NIH guidelines and were approved by UNC Animal Care and Use Committees (IACUC).

#### Single-nucleus RNA-seq of postnatal heart

*Library Preparation*

Hearts were harvested from mice at postnatal day 0 (p0, 2 replicates), day 7 (p7, 2 replicates), day 14 (p14, 2 replicates), and day 21 (p21, 2 replicates), respectively. Hearts were perfused with KCl and PBS before harvesting. After removing the heart from the chest, hearts were minced into pieces on a glass petri dish on ice using a razor blade. Minced heart tissue was then transferred to a 15 ml falcon tube for tissue homogenization with 1 ml of homogenization buffer (0.32M sucrose, 1mM Tris-HCl PH8.0, 3mM Magnesium Acetate, 0.1mM EDTA, 5mM Calcium Chloride, 0.1% Triton, 1x cOmplete Protease Inhibitor Cocktail, 200 U/ml RNase OUT, 1mM DTT). The tissue homogenization was performed on ice with two cycles of 30 second on and 60 sec off. After the tissue was completely homogenized, the tissue lysate was then transferred to a 7 ml tissue grinder. 1 ml of tissue homogenization buffer was used to wash the falcon tube before adding it to the tissue grinder. Eight strokes were performed with a loose pestle (A) on ice. 30 strokes were performed with a tight pestle (B) on ice. Then the tissue lysate was passed through a 70 µm and followed with a 40 µm cell strainer. Wash off any left-over tissue lysate from the 40 µm cell strainer with 1 ml of the homogenization buffer. Then the nuclei were pelleted by centrifuging at 500 g for 5 min at 4°C. Wash the pellet twice with resuspension buffer (PBS/1%BSA, 200 U/ml RNase OUT, 1 mM DTT). The nuclei pellets were then resuspended in 0.5 ml resuspension buffer. Before running the nuclei pellet with a 10x 3' scRNA-seq kit, isolated nuclei were labeled with DRAQ5 and sorted with flow cytometry to ensure the purity of the nuclei. The single-nucleus library preparations were performed strictly following the manufacturer's manual (Chromium Single Cell 3' Reagent Kits User Guide, CG000315).

*Preprocessing and cluster annotation*

Single-nucleus RNA sequencing data were processed and analyzed using the Seurat package (v5) in R. Raw count matrices were first filtered to remove low-quality cells based on mitochondrial gene expression (> 5%) and gene detection thresholds (nFeature > 7500 or nFeature <750). Then the doublet was identified and removed using the default setting of scDblFinder(*77*). Given that the single-nucleus libraries were prepared and synthesized in one batch, the single-nucleus RNA-seq libraries from different postnatal time points were directly integrated. Data normalization was performed using a regularized negative binomial regression method (SCTransform) to account for technical variability. Highly variable genes were then identified and used for dimensionality reduction via principal component analysis (PCA). IKAP was used to determine the number of clusters and the optimal principal component and cluster number pair(*33*)I. For visualization, Uniform Manifold Approximation and Projection (UMAP) was employed to project the high-dimensional data into two dimensions. Differential expression analysis between clusters was conducted using the Wilcoxon rank-sum test, and marker genes were identified based on an adjusted p-value cutoff of 0.05. The annotation of each cluster was determined based on the gene ontology enrichment analysis of the marker genes of each cluster and the published literature to ensure the annotation is biologically relevant.

*Prioritization of cell types responsive to maturation with Augur*

To identify cell types exhibiting the most pronounced transcriptional changes during maturation, we applied Augur (v1.0.3), a machine learning-based method for cell-type prioritization(*56*). Augur quantifies the separability of cell states across experimental conditions in high-dimensional gene expression space, assuming that cell types with stronger responses to a perturbation (here, maturation) will achieve higher separability. For each comparison, raw UMI count matrices with annotated cell types were used in Augur. We choose the default parameters, select the top 50% of highly variable genes, and randomly drop 50% of features in each run. Accounting for class imbalance, 20 cells per condition were randomly subsampled. The mean three-fold cross-validated area under the curve (AUC) was computed. Cell types were ranked by their mean AUC, where values approaching 1 signify near-perfect separation between conditions, and 0.5 represents random chance. We further perform the differential prioritizations to compare the difference between the time periods, e.g. P0-p7 versus p7-p14 and p7-p14 versus p14-p21.

*Weighted correlation network analysis of cardiomyocytes in the postnatal heart*

We subset snRNA-seq dataset to ventricle cardiomyocyte (v.CM) for WGCNA co-expression network analysis using hdWGCNA(*60*). Metacell transcriptomic profiles were constructed separately for each of the eight samples by aggregating transcriptome similar nuclei into one metacell using the hdWGCNA function MetacellsByGroups with following parameter: k=25, max_shared =10. We selected a soft-power threshold $\beta$=5 based on the parameter sweep performed with the TestSoftPowers function. The co-expression network was computed with the ConstructNetwork function with the following parameters: networkType = “signed”, TOMType = “signed”, soft_power = 5. Module eigengenes were computed using the ModuleEigengenes function. Eigengene-based connectivity for each gene was computed using ModuleConnectivity. The co-expression network was plotted with ModuleNetworkPlot with following parameter: n_inner=10, n_outer=20, n_conns=Inf. We used ClusterProlifer to perform gene ontology enrichment analysis on the top 200 genes in each module ranked by kME.

**Single-cell Spatial Transcriptome of postnatal heart**

*Gene panel design*

A panel of custom probes targeting 405 genes were built for xenium assay. 186 genes were selected as cell-type markers based on the snRNA-seq. 22 cardiomyocyte maturation-related genes were included. Additionally, 208 ligand and receptor genes were included based on the preliminary cell communication analysis on the snRNA-seq of the activated signaling pathways in the postnatal heart. The probe panels were fed into the Xenium panel designer together with the paired snRNA-seq library to ensure the panel design could provide enough resolution for cell type annotation in the spatial data.

*Xenium sample preparation and xenium instrument run*

Hearts were harvested from mice at postnatal day 0 (p0), day 7 (p7), day 14 (p14), and day 21 (p21), respectively. Hearts were perfused with KCl and PBS before harvesting. Then the heart was isolated from the mouse and transferred to cryomold with OCT. The tissue orientation was adjusted with forceps, and additional OCT was added to ensure the heart was fully submerged. Then, the cryomold with heart was lowered into precooled isopentane on dry ice without fully submerging. Once the OCT is frozen, the cryomold is placed on the aluminum foil on dry ice for 30 minutes before transferring to a -80 freezer for long-term storage. The 10 µm tissue sections of the postnatal heart for Xenium assay were sectioned using cryostat with a setting of -20°C for the blade and -10°C for the specimen. The adjacent tissue sections were collected to extract RNA for RNA quality control by measuring the DV200 using a Tape station. The Xenium slides were transported to Duke Molecular Genomics Core for Xenium Run with Xenium Cell Segmentation Staining Kit. The TA variety of output files was produced by the on-instrument pipeline. The essential files used downstream were the feature-cell matrix (HDF5 and MEX formats identical to those output by Xenium Ranger), the transcripts (listing each mRNA, its 3D coordinates, and a quality score), and the cell boundaries CSV file. These files were then transferred for downstream analysis off-instrument.

*Spatial Resolution single-cell regulatory network inference Analysis*

To construct transcriptional regulatory networks during maturation, we employed the SCENIC workflow (version 1.3.1) on snRNA data(*51*). SCENIC utilizes single-cell gene expression data to infer regulons, defined as transcription factors (TFs) and their target genes. First, the co-expression network was inferred using GRNBoost2(*78*), identifying genes co-expressed with TFs and separating targets into positively and negatively correlated groups. Genes with low expression levels or low positive rates were filtered using the geneFiltering function with default parameters. Next, RCisTarget (version 1.10.0) performed cis-regulatory motif analysis using the cisTarget databases (mm10 refseq_r80 v9 database) to select significantly enriched motifs among the co-expression modules. The TF-gene modules and target gene predictions were integrated to construct regulons. Finally, AUCell (version 1.20.2) was used to score regulon activity in each cell by calculating the AUC based on gene expression rankings. The identified TFs were integrated onto spatial location by Cytospace (version 0.0.3) and visualized using the pheatmap package (version 1.0.12) to compare the difference across cell types and time points.

*Spatial Clustering of postnatal heart with CellCharter*

To identify the cellular neighborhoods within the postnatal heart across the postnatal development, we applied CellCharter to our Xenium datasets. Briefly, the count matrix and the spatial coordinates of each cell were loaded using scanpy into python environment. Then, the dimensionality of the transcriptome data was reduced using scVI with the default parameter before computing the spatial cluster. To connect each cell as a network for spatial clustering, squidpy's gr.spatial_neighbors function was used, followed by gr.remove_long_links to remove the connection between distant cells. Then, cellcharter's function cc.gr.aggregate_neighbour was used to perform neighbourhood aggregation for each cell. To determine the optimal cluster number, cc.tl.ClusterAutoK was performed to obtain the best candidates for the number of spatial clusters. Finally, autoK.predict was performed with k=12 to assign the cellular neighborhoods to each cell in the postnatal heart.

*Maturation index and cellular neighborhood analysis*

To infer the maturation status of each cardiomyocyte in the postnatal heart, we calculated the maturation index based on the expression of the 22 maturation-related genes from the custom xenium panels. The maturation index is calculated with AddModuleScore function from Seurat v5. Next, to infer the correlation of the maturation of cardiomyocyte to its relative position of other non-CM cardiac cells, sq.gr.spatial_neighbors from squidpy(*79*) was used to build a spatial graph and the adjacency matrix of each sample. Then, based on the spatial adjacency matrix, each cell's neighbor level was assigned from 0 to 10 based on the ordinal numbers of hexagonal rings the cell resides in, where 0 is the cell itself, and 10 is the $10^{th}$ rings and above. Pearson correlations were performed to test the correlation between the maturation index and the cellular neighborhood level for each cell type accordingly.

*Embedding single-nucleus transcriptomes in space by Cytospace*

To spatially resolve transcriptional states across the developing mouse neonatal heart, we embedded snRNA-seq data into a common coordinate framework using Cytospace (version 1.1.0)(*32*). This approach integrates matched time point snRNA-seq and Xenium datasets by optimizing the spatial assignment of single-cell barcodes while preserving their transcriptional similarity and spatial coherence. We provided Cytospace with (i) the snRNA-seq count matrix (cell type annotations included) and (ii) the ST count matrix with spatial coordinates. Cytospace's optimal transport algorithm was run with default parameters, minimizing a cost function balancing transcriptional similarity (snRNA-seq-to-ST gene expression correlation) and spatial smoothness (distance between neighboring spots). Mapping accuracy was assessed by comparing Cytospace-assigned cell types to manual annotations in ST data.

*Cell–cell communication*

Cell–cell communication in spatial data was assessed using COMMOT (version 0.0.3)(*41*), which accounts for the competition between different ligand and receptor species as well as spatial distances between cells. We used CellPhoneDB_v4.0 ligand-receptor database by commot.pp.ligand_receptor_database() function. Secreted signaling: L-R pairs (e.g., VEGFA–FLT1) were inferred with a maximum allowed spatial distance of 200 µm, reflecting the diffusion range of soluble ligands. ECM-receptor interactions (e.g., FN1–ITGA5) and cell-cell contact pairs (e.g., CDH1–CDH1) were restricted to 50 µm, consistent with direct membrane proximity. The commot.tl.spatial_communication function was run with heteromeric=True to include multi-subunit LR complexes (e.g., IL2 receptor (IL2RA–IL2RB–IL2RG)).

*Signaling-to-transcription regulatory NETwork (SigNET)*

To identify transcription factors (TFs) potentially regulated by spatially resolved ligand-receptor interactions, we integrated COMMOT-derived L-R interaction scores with single-cell TF activity scores inferred by SCENIC (version 1.3.1) using arboreto package (version 0.1.6) with function GRNBoost2(*78*), a gradient-boosting machine-based framework for gene regulatory network inference.

With COMMOT inferred ligand-receptor expression score and SCENIC inferred transcription factor score, we utilized GRNBoost2, a gradient boosting machine-based GRN inference algorithm, to compute the co-expression module. Top TF with 1 standard deviation (1 SD) above mean weight were selected. This approach integrates ligand-receptor interactions into the GRN framework, enabling the identification of regulatory modules that may drive biological processes.

*Machine Learning Prediction on Maturation Index*

To find the important features that are associated with the maturation index, we use Random Forest regression pipeline with SHAP (SHapley Additive exPlanations, version 0.44.1) to find the most important features. We use scikit-learn (version 1.3.2) package for pipeline building, using

RandomForestRegressor() function with 100 trees. The model performance was evaluated using $R^2$ (coefficient of determination) on the test set. To interpret feature importance, SHAP values were computed using TreeExplainer with the tree_path_dependent approximation method. A subset of 300 samples from the test set was used for SHAP calculations to optimize runtime performance.

**in vivo Probe-based Indel-detectable Perturb-seq (PIP-seq)**

*AAV9 CRISPR Guide Library Design and AAV9 Preparation*

Two sgRNAs targeting at candidate genes were designed using CRISPick (https://portals.broadinstitute.org/gppx/crispick/public), selecting Mouse GRCm38 as the reference genome and CRISPRko as mechanism. Top 2 sgRNAs based on the overall on-target and off-target rank were selected for cloning. Two sgRNAs were cloned sequentially to the CASAAV backbone following the published protocol(*73*). Sanger sequencing was performed after each cloning to ensure the correct insertion of sgRNA. To produce the AAV9 CRISPR guide library, the 53 sgRNA contained plasmids were miniprepped and pooled in equal molar before being sent to the UNC Vector Core for AAV9 production.

*sgRNA detection and perturbation detection probe design*

Probes for the sgRNAs detection and perturbation detection were obtained from IDT as oPools at 10 pmol/oligo. Probe sequences are found in the Supplementary Table 3 and Table 4.

*PIP-seq library preparation*

AAV9 library was injected into neonatal mice at postnatal day 0. Hearts were harvested on postnatal day 14. Hearts were perfused with KCl and PBS before harvesting. After removing the heart from the chest, the ventricles of the harvested heart were minced into pieces on a glass petri dish on ice using a razor blade. A cut/bore tip was used to transfer tissue pieces from the glass petri dish to a 15ml falcon tube. The tube was then topped with fixation buffer (4% formaldehyde, 1X Fix & Perm Buffer) to reach a 25 mg tissue to 1ml fixation ratio. The tissue pieces were then incubated in the fixation buffer for 20 hours at 4°C without rotation. The incubation time should be consistent between samples for the same batch of experiments. After incubation, tissue pieces were centrifuged at 850 g for 5 min at room temperature. The supernatant was removed without disturbing the tissue pellet. Then the tissue pieces were washed again with 2 ml chilled PBS and resuspended in tissue homogenization buffer for nuclei isolation. After nuclei isolation, flow cytometry was performed to enrich the GFP+/DRAQ+ population. Then, fixed RNA single-nucleus libraries were generated with the GFP+/DRAQ5+ population by strictly following the manufacturer's manual (Chromium Fixed RNA Profiling Reagent Kits for Singleplexed Samples, CG000691).

*In vivo Perturb-seq analysis*

To map sequencing read to the mouse transcriptome and our sgRNA sequence, a customized csv file was generated containing the probe sequences of the whole mouse transcriptome and sgRNAs. Cellranger aggr function was used to generate the count matrix. To access the perturbation effect of the sgRNAs on the cardiomyocyte transcriptome, we calculated the module score for gene sets related to the cardiomyocyte maturation process.

$$[\sum_{g \in module} \frac{x_{gc} - \mu_{gp}}{\sigma_{gp}}] \times \frac{1}{N_{modules}}$$

To identify the sgRNA showing significant change to each module, Wilcoxon Rank Sum Test was performed between the perturbed nuclei and non-targeting nuclei. To access the differentially expressed genes in perturbed nuclei, Findmarker function from Seurat package was used between the perturbed nuclei population and non-targeting nuclei population.

***Animal experiments***

*AAV9 injection and Echocardiogram*

AAV9 library was injected into neonatal mice at postnatal day 0 as a high-dosage treatment. Echocardiography was performed on a VisualSonics Vevo 2100 with Vevostrain software. Animals were awake during this procedure and held in a standard handgrip. Standard short-axis M-mode measurements were recorded. Echocardiography was performed blinded to the treatment group. At the end of the study, standard morphometric measures were obtained, including body and heart weights and tibia length.

*Cardiomyocyte isolation and immunocytochemistry*

Cardiomyocyte isolation was carried out as previously described with minor modifications(*81*). Briefly, Male mice aged 4 weeks were anesthetized, and their chests were surgically opened to expose the heart. After cutting the descending aorta, the heart was immediately perfused with 7 ml of EDTA buffer via the right ventricle. Tissue digestion was initiated by sequentially perfusing the left ventricle (LV) with EDTA buffer, followed by perfusion buffer and collagenase buffer. The ventricles were then dissected into approximately 1 mm fragments using forceps. Cellular dissociation was completed by the addition of 5 ml of stop buffer and then was passed through a 100 μm filter. 3 sequential rounds of gravity settling were required to form a highly pure myocyte fraction. For immunocytochemistry, the isolated cardiomyocytes were fixed with 4% PFA at room temperature for 15 minutes and permeabilized with 0.2% Triton for 15 minutes. After blocking with 5% BSA, fixed cells were stained with primary antibody. The primary antibody used was anti-α-Actinin mouse monoclonal antibody (Sigma, #A7811, 1:400). Cell nuclei were counterstained with Hoechst 33342 at 1 μg/ml. Cell size was measured and quantified by using ImageJ software.

*Immunostaining and immunofluorescence analysis*

Mouse hearts were fixed with 0.5% PFA overnight at 4°C and embedded with OCT after dehydration. The myocardial tissues were serially sectioned at a thickness of 10 µm. Then, sections were washed twice with 0.1% PBS-Tween for 5min and permeabilized with 0.2% PBS-Triton for 15min. 5% BSA was used to block the sections for 1 hour. After that, the sections were incubated with Alexa Fluor-555 conjugated WGA antibody (ThermoFisher Scientific) for 1 hour at room temperature. All slides were mounted and analyzed with EVOS FL cell imaging system. The cross-sectional area of the cardiomyocytes outlined by WGA staining was measured and quantified by using ImageJ software.

*In situ confocal imaging*

*In situ* T-tubule imaging was performed following a published protocol(*82*). In brief, hearts were dissected from euthanized mice and first perfused with PBS. Perfusion buffer containing 100 µg/ml FM 4–64 (Invitrogen, 13320) was loaded into the heart by retrograde perfusion at room temperature for 10 min. The heart was next removed from the perfusion system, positioned on a glass-bottom dish, and immediately imaged by confocal microscopy (Zeiss 880).

### Supplementary Figures

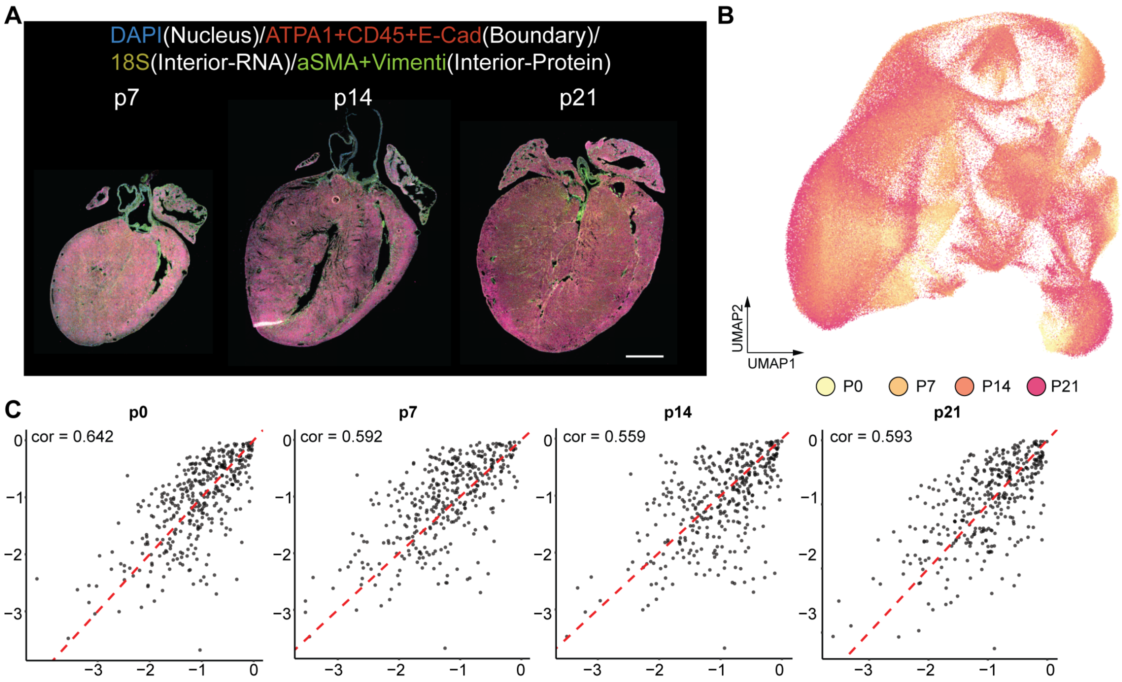


**Figure S1. Postnatal heart atlas with spatial and temporal resolution**

A) Multimodal segmentation enables improved segmentation of the multinucleated cardiomyocyte (indicated by the asterisk) in the postnatal hearts from various time points (p7, p14, and p21). Scale bar, 800 µm.

B) Approximately 403,584 cardiac cells's spatial transcriptome were profiled and clustered into distinct cell populations as shown in the UMAP and across four postnatal timepoints (colored accordingly).
C) Pearson correlation value between the single-nucleus RNA-seq and Xenium for each time point.

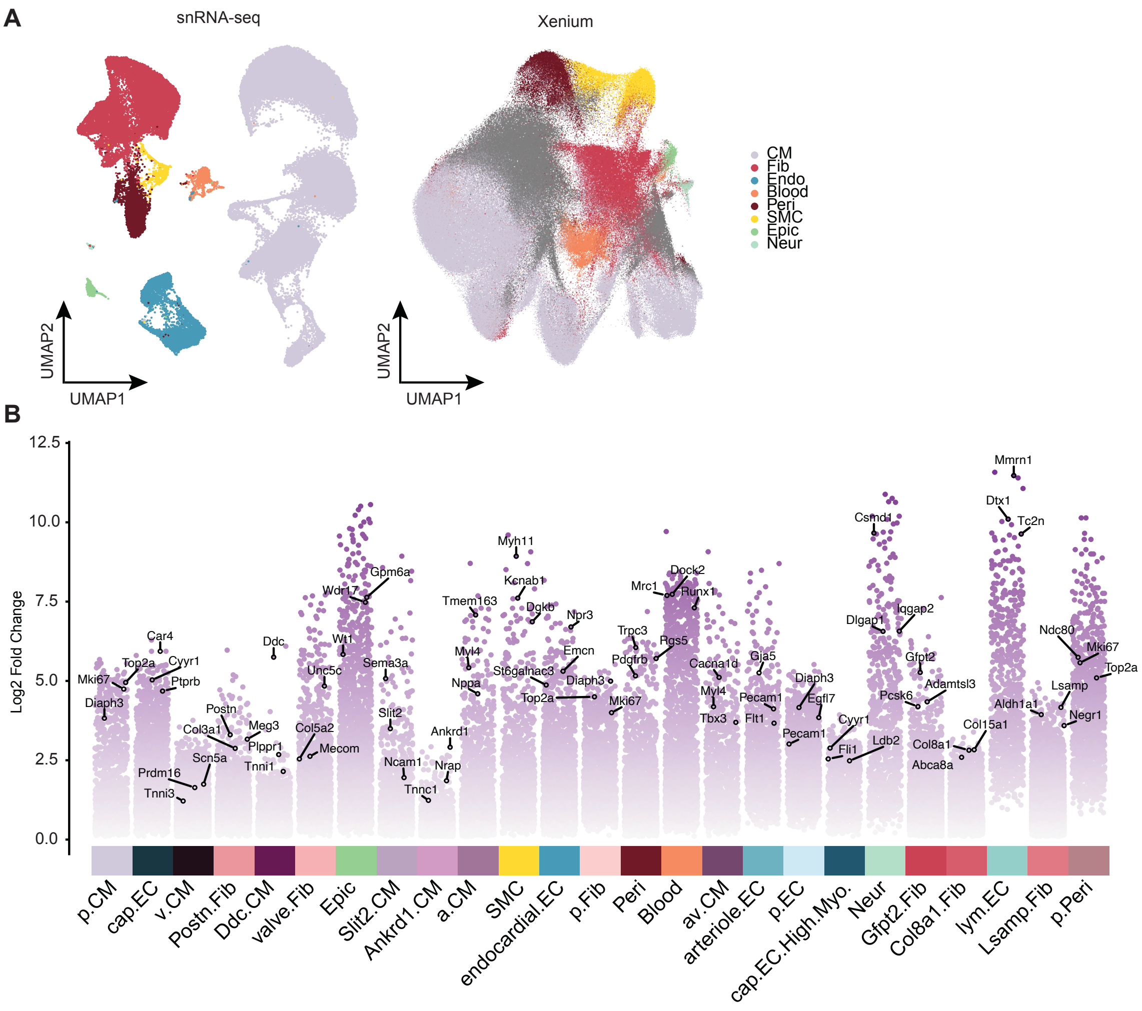


**Figure S2. Annotation of cell types in the postnatal heart atlas**

A) Nine cell types colored in the UMAP of snRNA-seq(Left) and Xenium (Right).

B) The marker genes for each of the 25 subordinate cell states. Selected top marker genes for each cell state were labelled. Y-axis is the $Log_2$(Fold Change) of expression between cell state to the remaining cardiac cells.

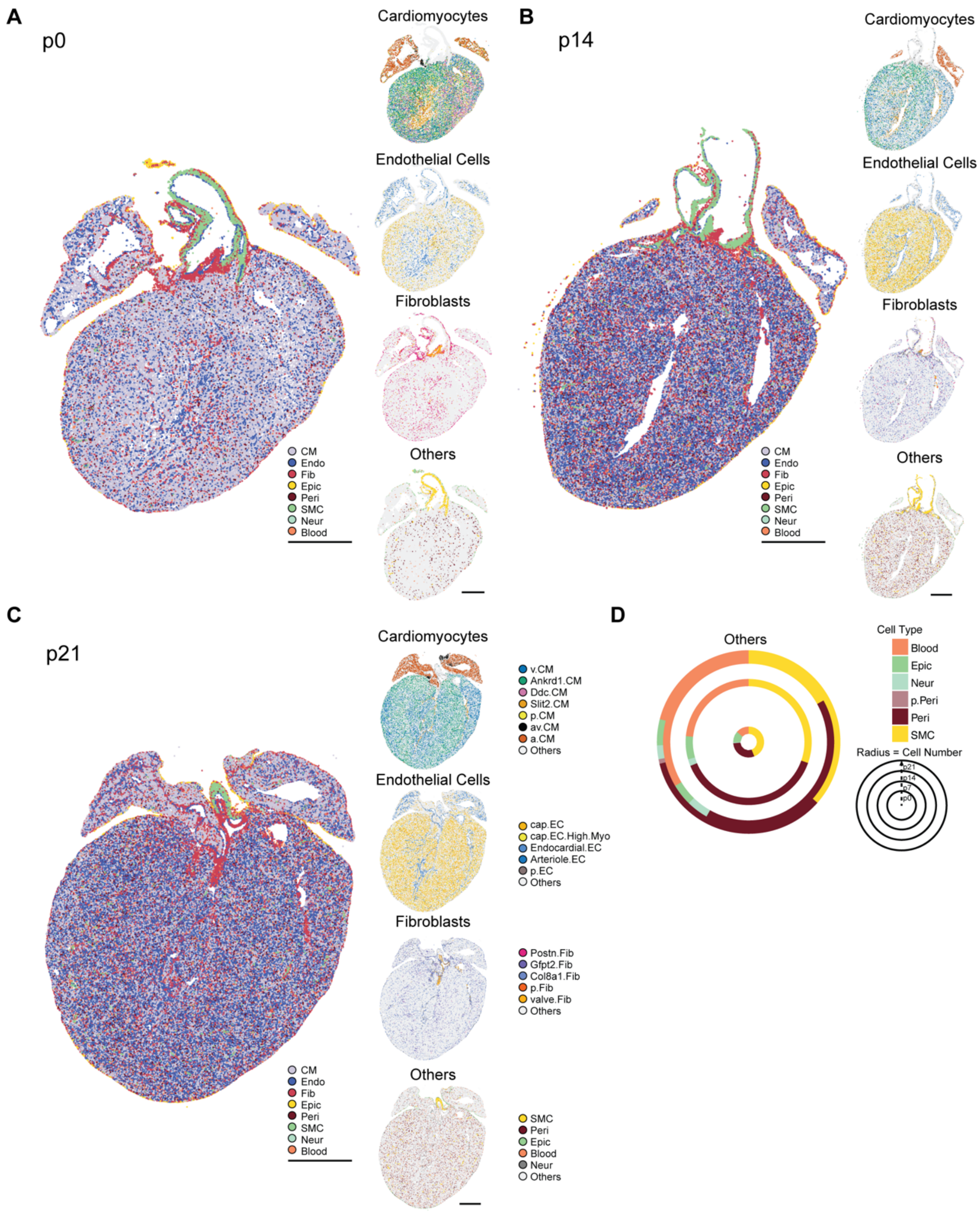


**Figure S3. Spatial location of the cell states in the postnatal hearts**

A) Left: Spatial map of heart at postnatal day 0. Each cell type was labeled with different color. Right: Spatial map showing the subordinate cell states of cardiomyocytes, endothelial cells, fibroblasts, and others in the heart at postnatal day 0. Scale bar, 500µm.

B) Left: Spatial map of heart at postnatal day 14. Each cell type was labeled with different color. Right: Spatial map showing the subordinate cell states of cardiomyocytes, endothelial cells, fibroblasts, and others in the heart at postnatal day 14. Scale bar, 1000µm.

C) Left: Spatial map of heart at postnatal day 21. Each cell type was labeled with different color. Right: Spatial map showing the subordinate cell states of cardiomyocytes, endothelial cells, fibroblasts, and others in the heart at postnatal day 21. Scale bar, 800μm.
D) Donut plot showing the changes in the proportion of remaining cell states (other than CM, EC, and Fib) in the postnatal heart over the postnatal development. The total cell number of each time point is correlated with the radius of the donut.

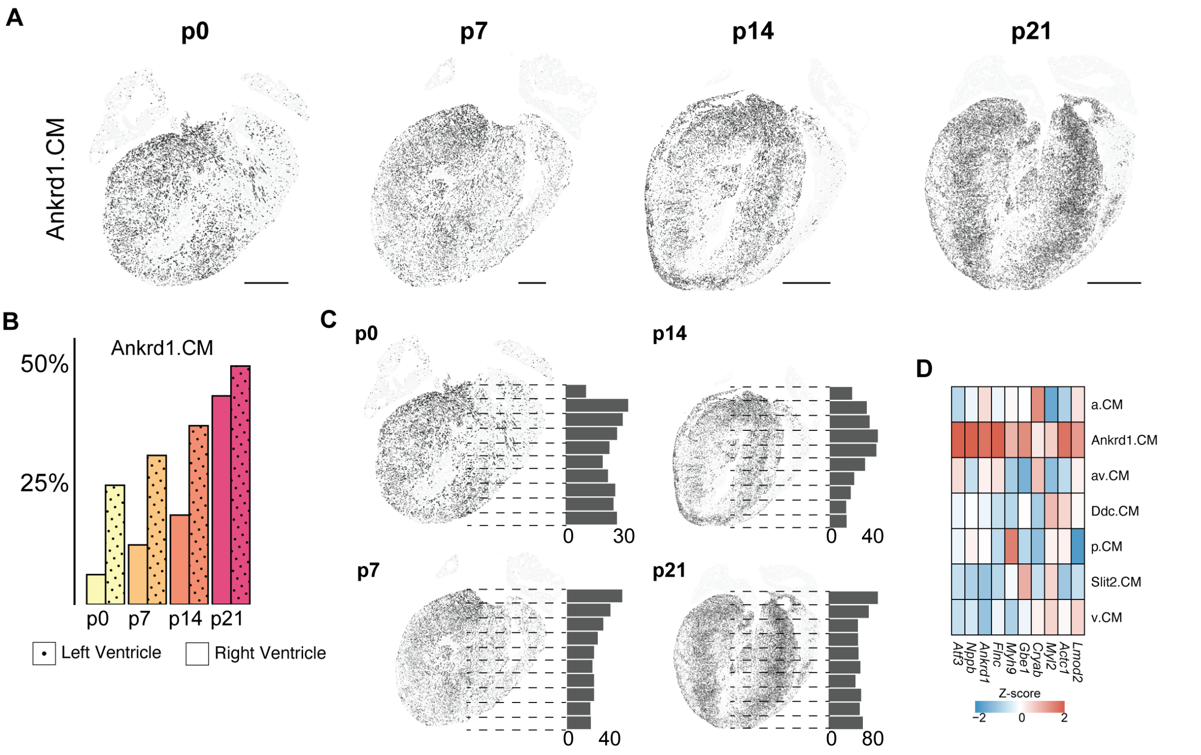


**Figure S4. Ankrd1.CM in the postnatal heart**

A） Spatial localization of the Ankrd1.CM in the postnatal hearts. Scale bar (p0, p7), 500μm; Scale bar (p14, p21), 1000μm.

B） Bar chart showing the percentage of Ankrd1.CM over all ventricle CMs in the left (Dot) and right (Plain) ventricle throughout the postnatal development.

C） The distribution of Ankrd1.CM along the vertical long axis of the postnatal heart. The vertical long axis is divided into 10 bins, and the percentage of Ankrd1.CM in each ventricle's CMs is represented as a bar plot next to the spatial localization images of the corresponding postnatal hearts.

D） Heatmap showing the expression of up-regulated genes in Ankrd1.CM compared to the other subordinate CM states in the postnatal heart. Color scheme represents the Z-score of the expression level from high (red) to low (blue).

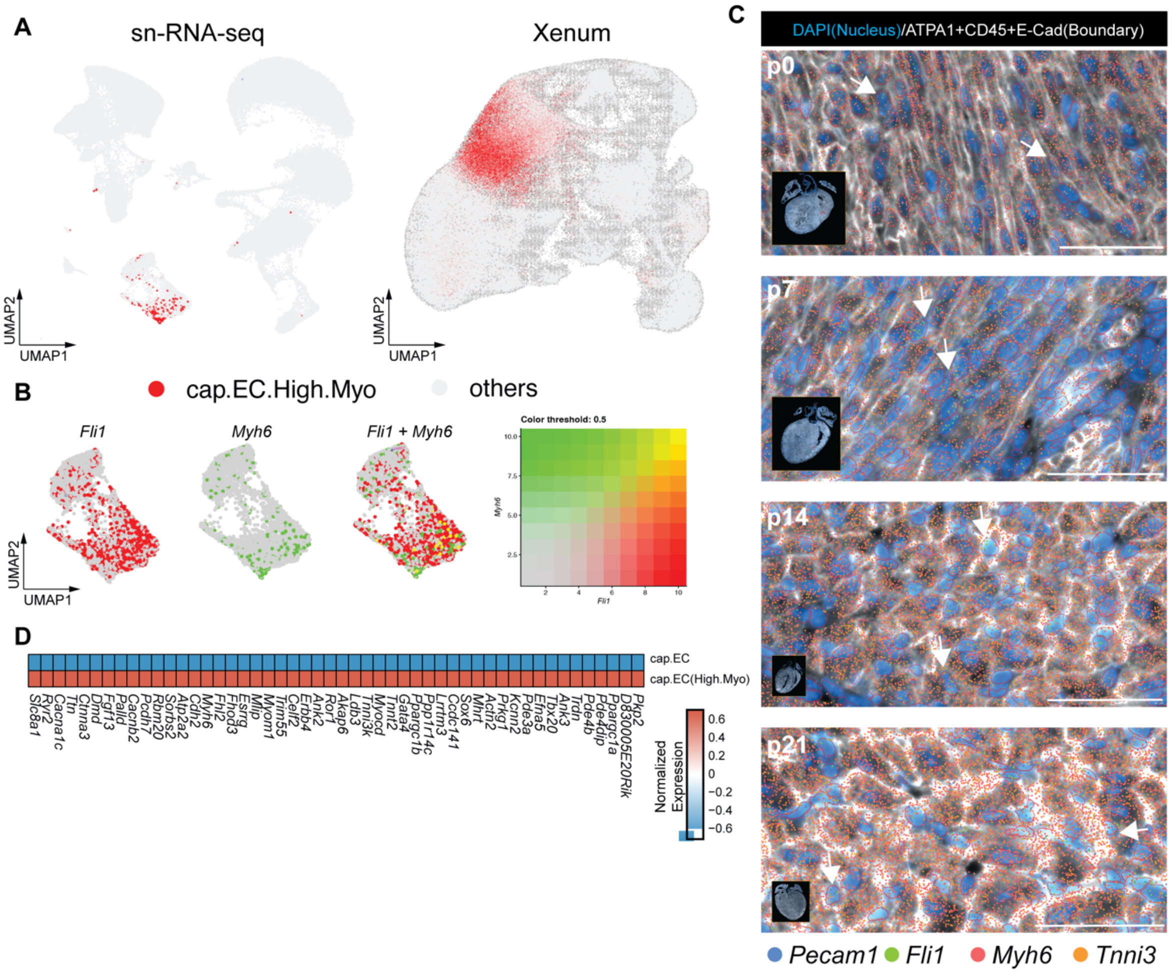


**Figure S5. Capillary ECs with high myocyte gene expression in the postnatal heart**

A) Cap.EC.High.Myo is presented in both the snRNA-seq and Xenium of the postnatal heart as highlighted as red dot in the UMAP of both datasets.

B) Co-expression of Endothelial Cells marker gene, Fli1, and cardiomyocyte marker gene, Myh6, within the same cell based on the snRNA-seq. Yellow indicates the co-expression of both genes, while green indicates Myh6-specific expression and red indicates Fli1-specific expression in the cell.

C) Co-expression of Endothelial Cells marker gene, Fli1, and cardiomyocyte marker gene, Myh6, within the same cell based on the Xenium. *Pecam1*, and *Fli1* are endothelial cell markers, while *Myh6* and *Tnni3* are cardiomyocyte cell markers. Scale bar, 25 µm.

D) Heatmap showing the expression of up-regulated genes in Cap.EC.high.myo compared to the cap.EC state in the postnatal heart. Color scheme represents the Z-score of the expression level from high (red) to low (blue).

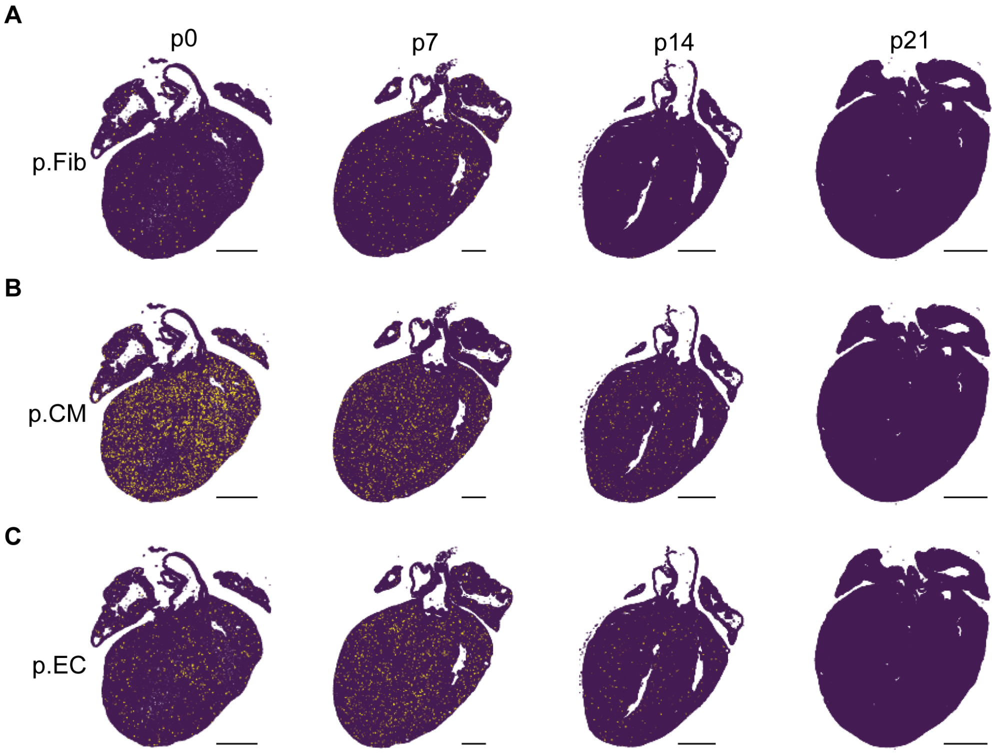


**Figure S6. Proliferative cell states in the postnatal heart**

A) Spatial map of proliferative fibroblast throughout the postnatal development. Scale bar (p0, p7), 500μm; Scale bar (p14, p21), 1000μm.

B) Spatial map of proliferative cardiomyocyte throughout the postnatal development. Scale bar (p0, p7), 500μm; Scale bar (p14, p21), 1000μm.

C) Spatial map of proliferative endothelial cells throughout the postnatal development. Scale bar (p0, p7), 500μm; Scale bar (p14, p21), 1000μm.

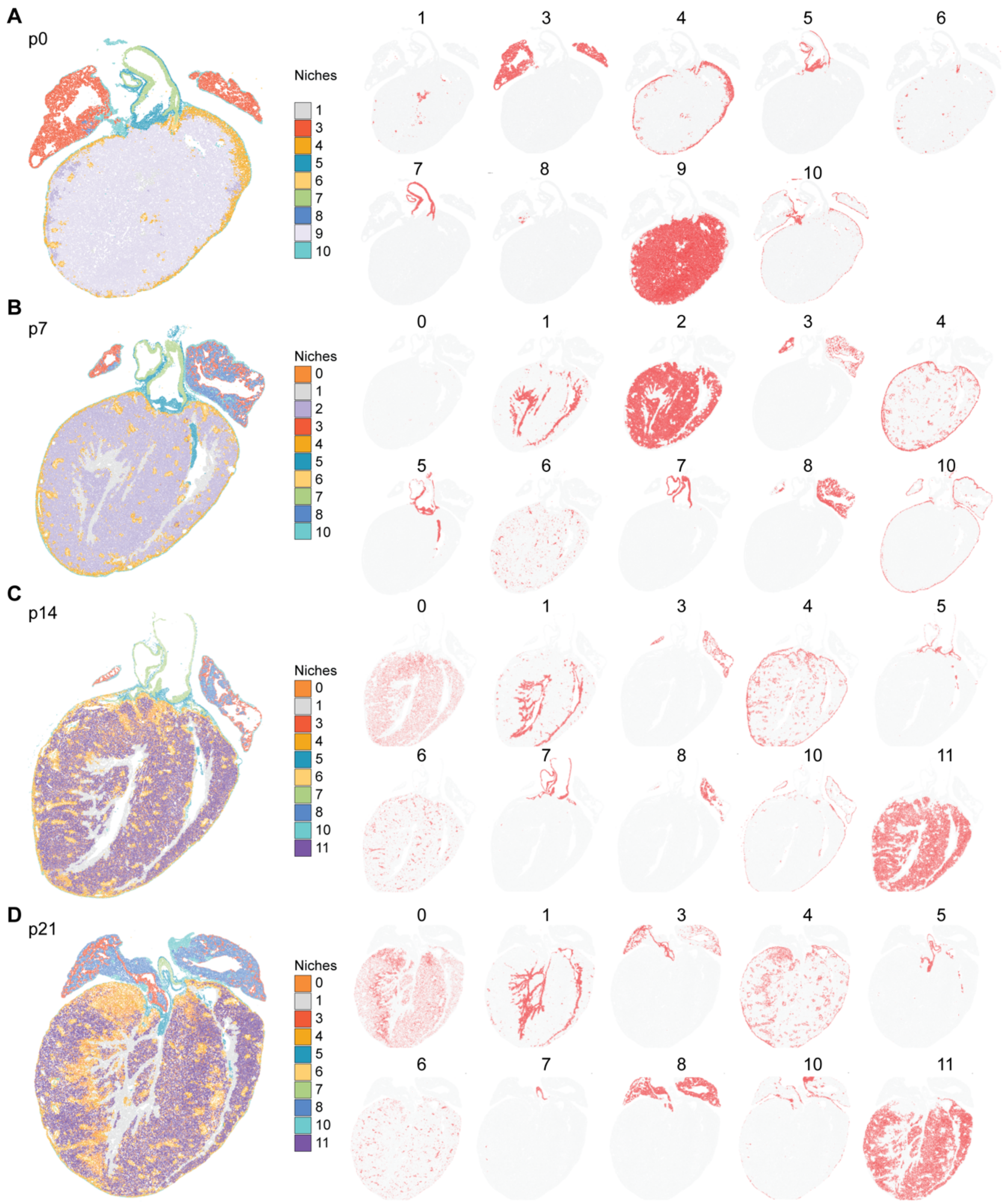


**Figure S7. Cell neighborhoods (niches) in the postnatal heart**

A) Left: Spatial map of heart at postnatal day 0. Each niche was labeled with different colors. Right: Spatial map of each niche in the postnatal heart at day 0. Scale bar, 500µm.

B) Left: Spatial map of heart at postnatal day 7. Each niche was labeled with different colors. Right: Spatial map of each niche in the postnatal heart at day 7. Scale bar, 500µm.

C) Left: Spatial map of heart at postnatal day 14. Each niche was labeled with different colors. Right: Spatial map of each niche in the postnatal heart at day 14. Scale bar, 1000µm.

D) Left: Spatial map of heart at postnatal day 14. Each niche was labeled with different colors. Right: Spatial map of each niche in the postnatal heart at day 14. Scale bar, 1000µm.

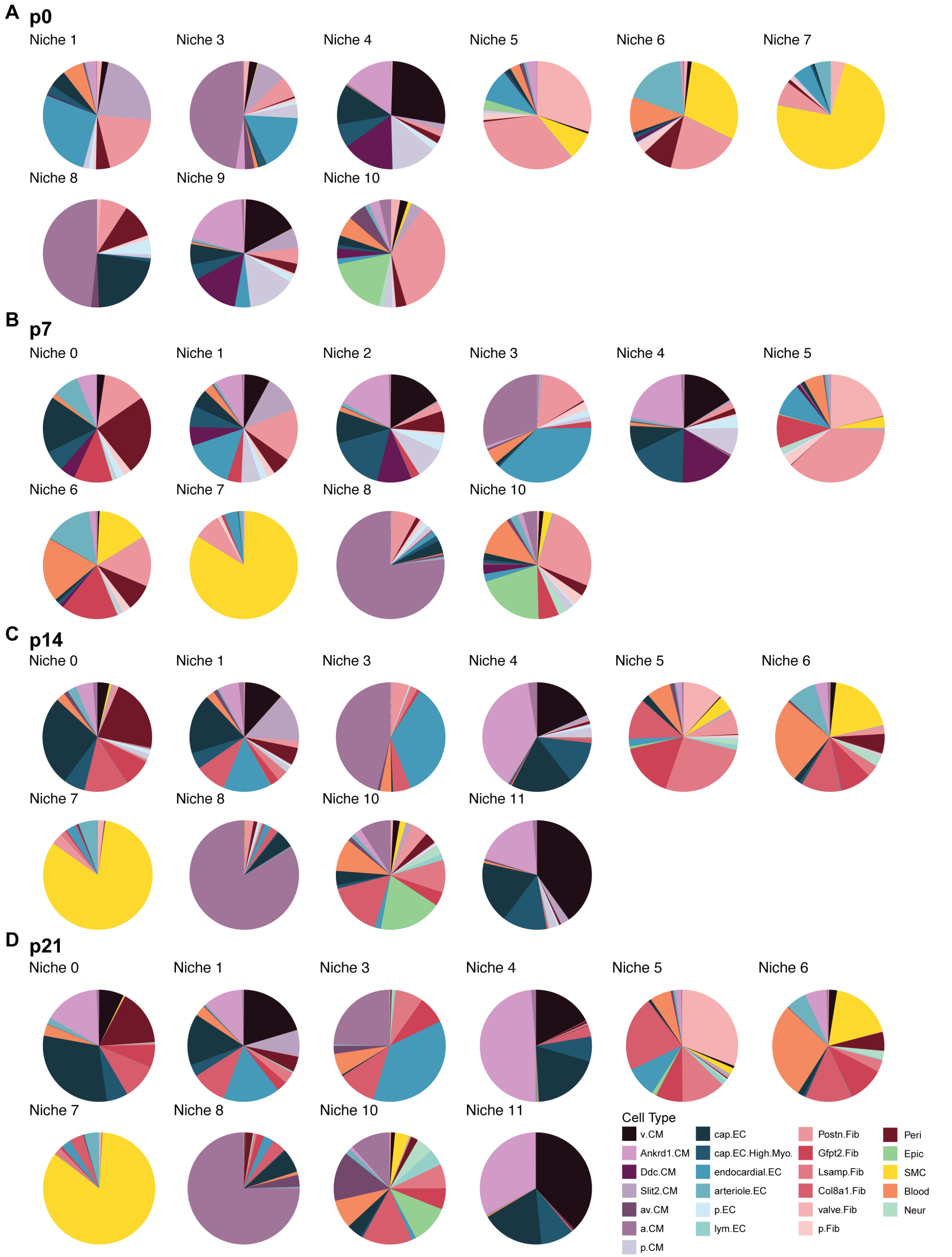


**Figure S8. Cell proportion of each niche in the postnatal heart**

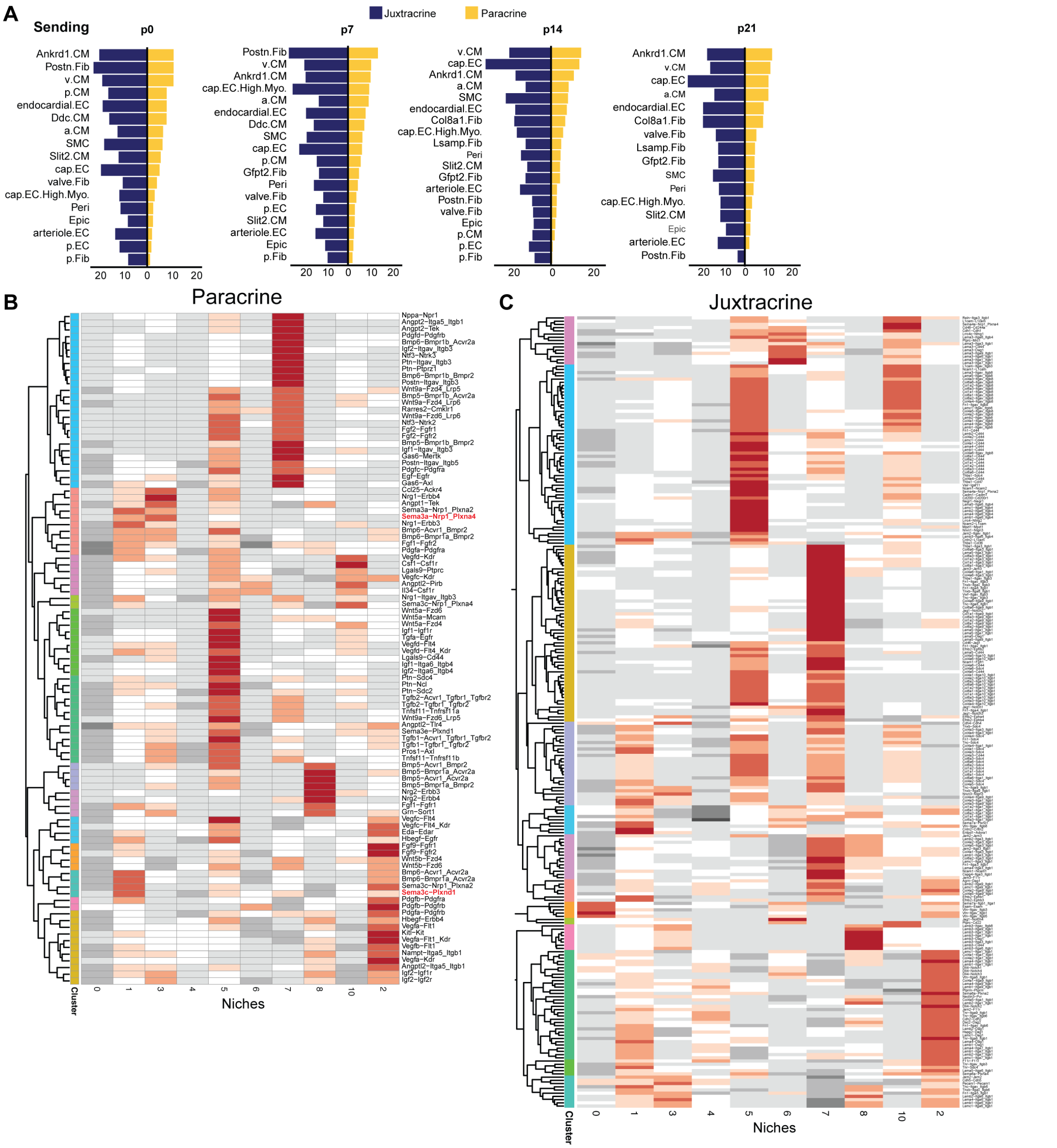


**Figure S9. Cell communications in the postnatal hearts**

A) The paracrine (Yellow) and juxtracrine (Naval blue) signaling intensity sent out by each cell type in the postnatal heart.

B) Heatmap showing the paracrine ligand-receptor pairs' activity in each niche in the postnatal heart at day 7.

C) Heatmap showing the juxtracrine ligand-receptor pairs' activity in each niche in the postnatal heart at day 7.

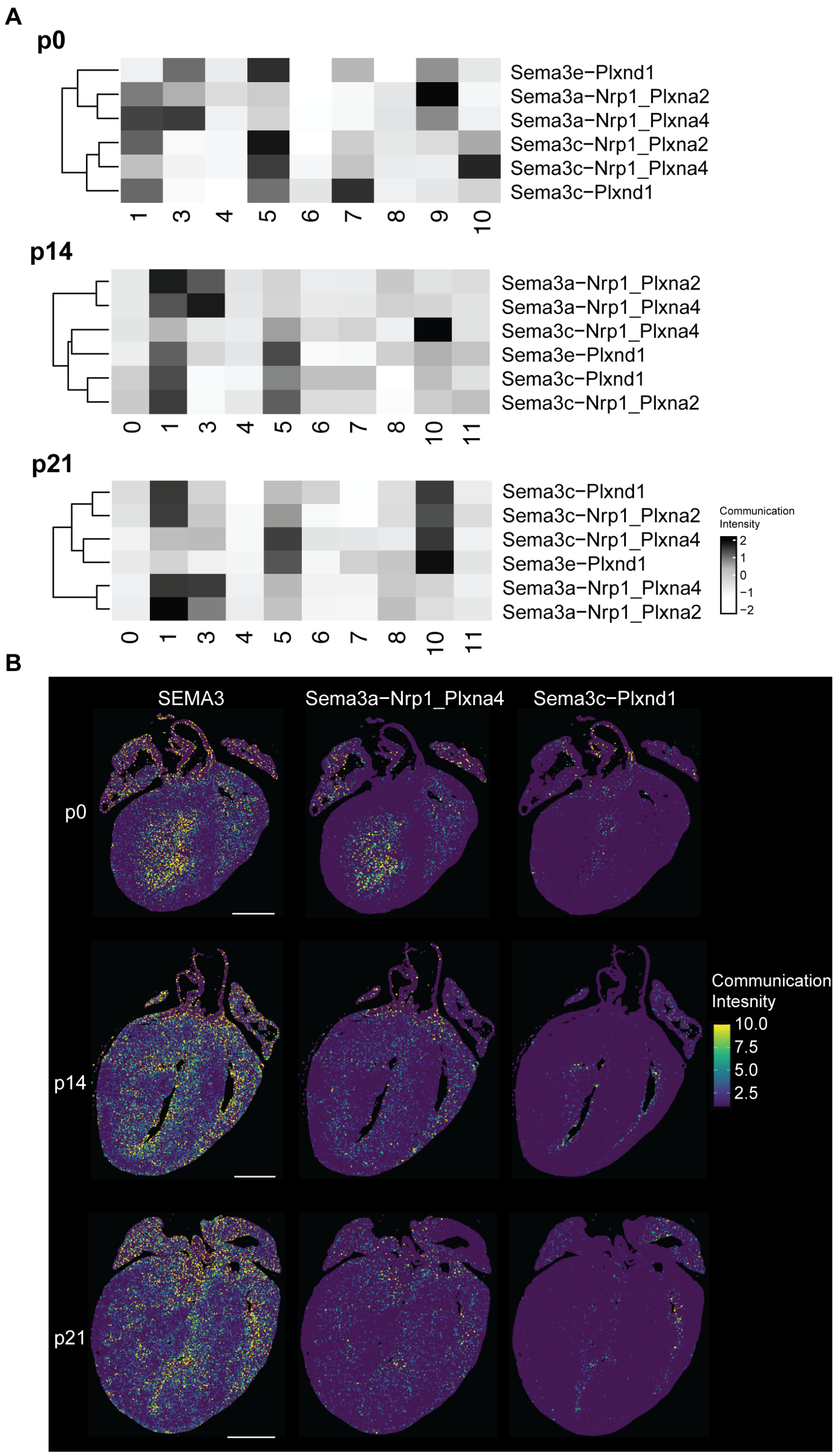


**Figure S10. SEMA3 signaling pathways in the postnatal hearts**

A) Heatmap showing the relative communication intensity of each ligand-receptor pair of the SEMA3 pathway at different cellular neighborhoods of the postnatal day 0, day 14, and day 21 heart.

B) The spatially resolved activities of the SEMA3 pathway and two ligand-receptor pairs, Sema3a-Nrp1_Plxna4 and Sema3c-Plxnd1 at postnatal day 0, day7, and day 21 heart. The white dashed area highlights the differential spatial activities of two ligand-receptor pairs. Scale bar (p0), 500µm; Scale bar (p14, p21), 1000µm.

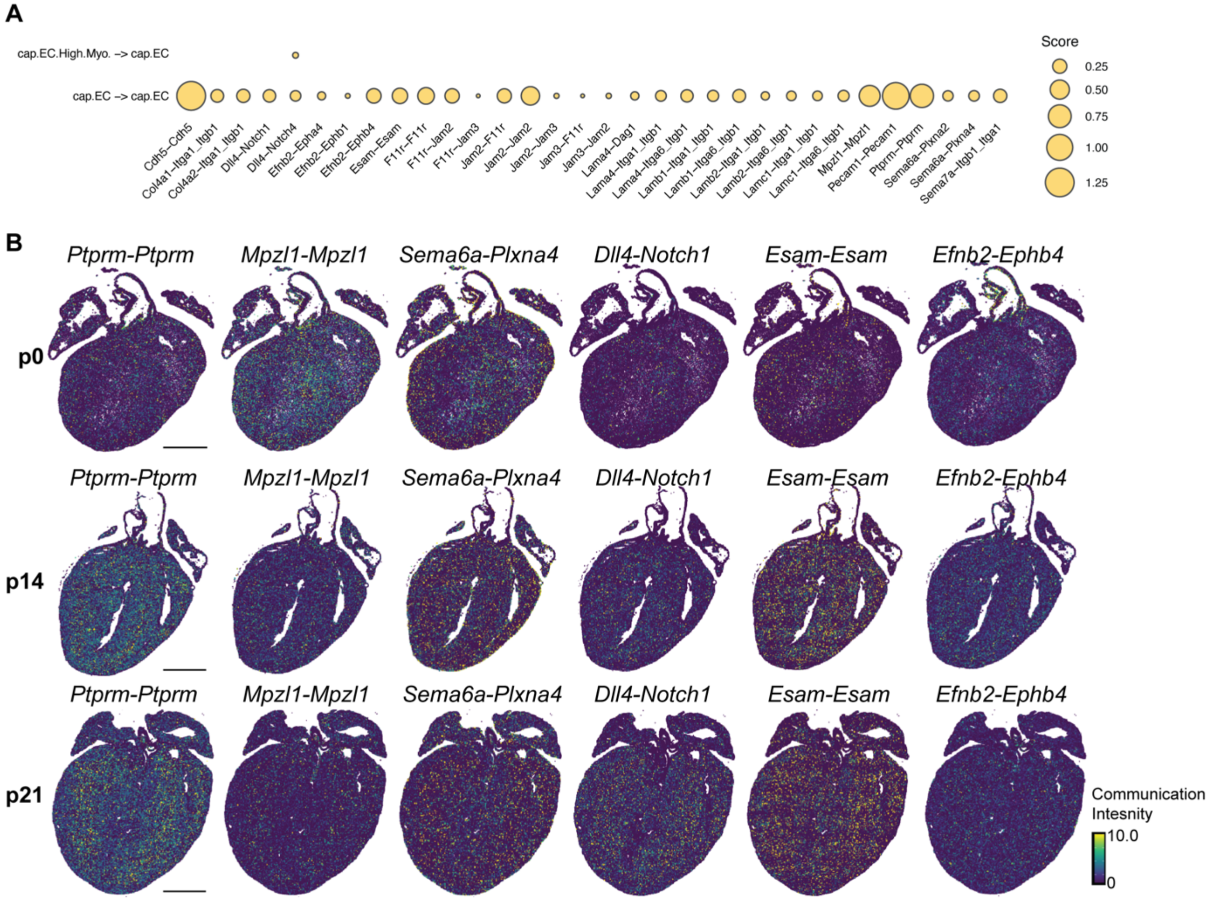


**Figure S11. SEMA3 signaling pathways in the postnatal hearts**

A) Dot plot showing the activated juxtracrine ligand-receptor pairs received by the cap.EC in the ventricle nicheThe size of each dot represents the normalized communication intensity of each ligand-receptor pair.

B) The spatially resolved activities of *Efnb2-Ephb4*, *Esam-Esam*, *Sema6a-Plxna4*, *Dll4-Notch1*, *Ptprm-Ptprm*, and *Mpzl1-Mpzl1* at postnatal day 0, day 14, and day 21 heart. The color scheme represents the normalized communication intensity.

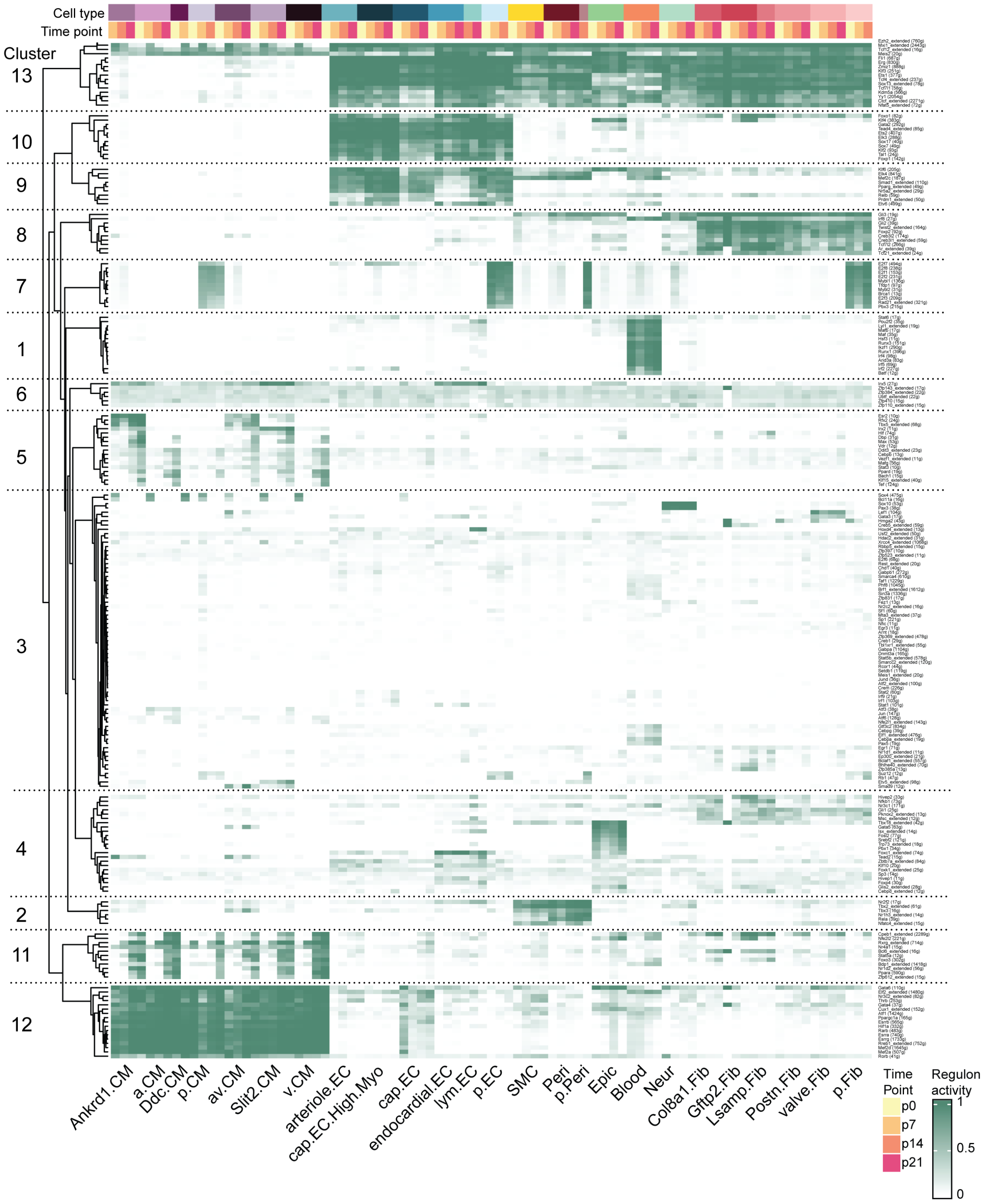


**Figure S12. SEMA3 signaling pathways in the postnatal hearts**

Heatmap showing unsupervised hierarchical clustering of the regulon activity in each cell type at each postnatal time point. The color Scheme represents the regulon activity from high (Green) to low (White).

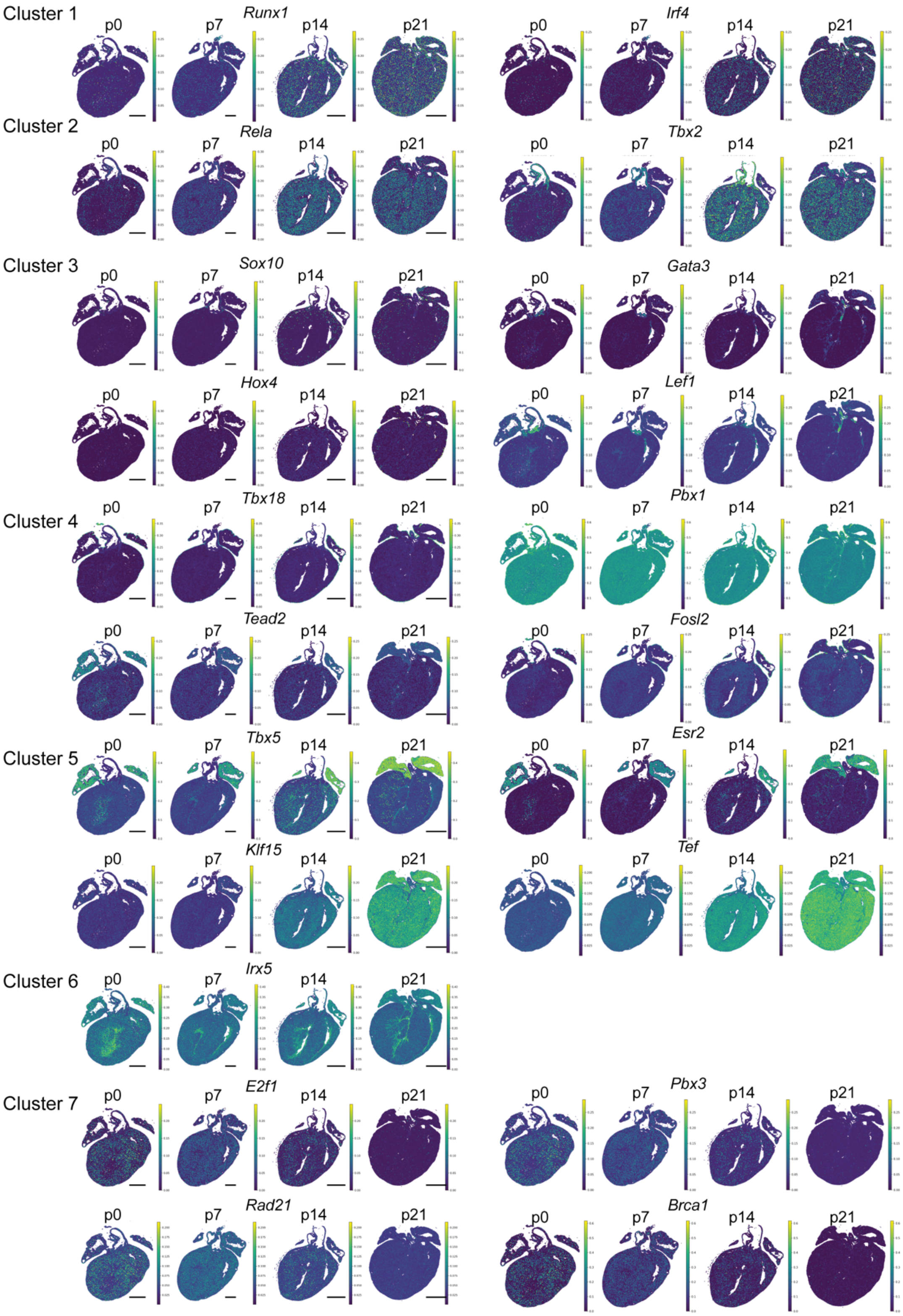


**Figure S13. Regulon activities in the postnatal development for cluster 1 to cluster 7**

Representative regulon activities were displayed in the postnatal hearts spatially and temporally. The color scheme represents the regulon activity of each cell in the postnatal heart from high (green) to low (blue). Scale bar (p0, p7), 500µm; Scale bar (p14, p21), 1000µm.

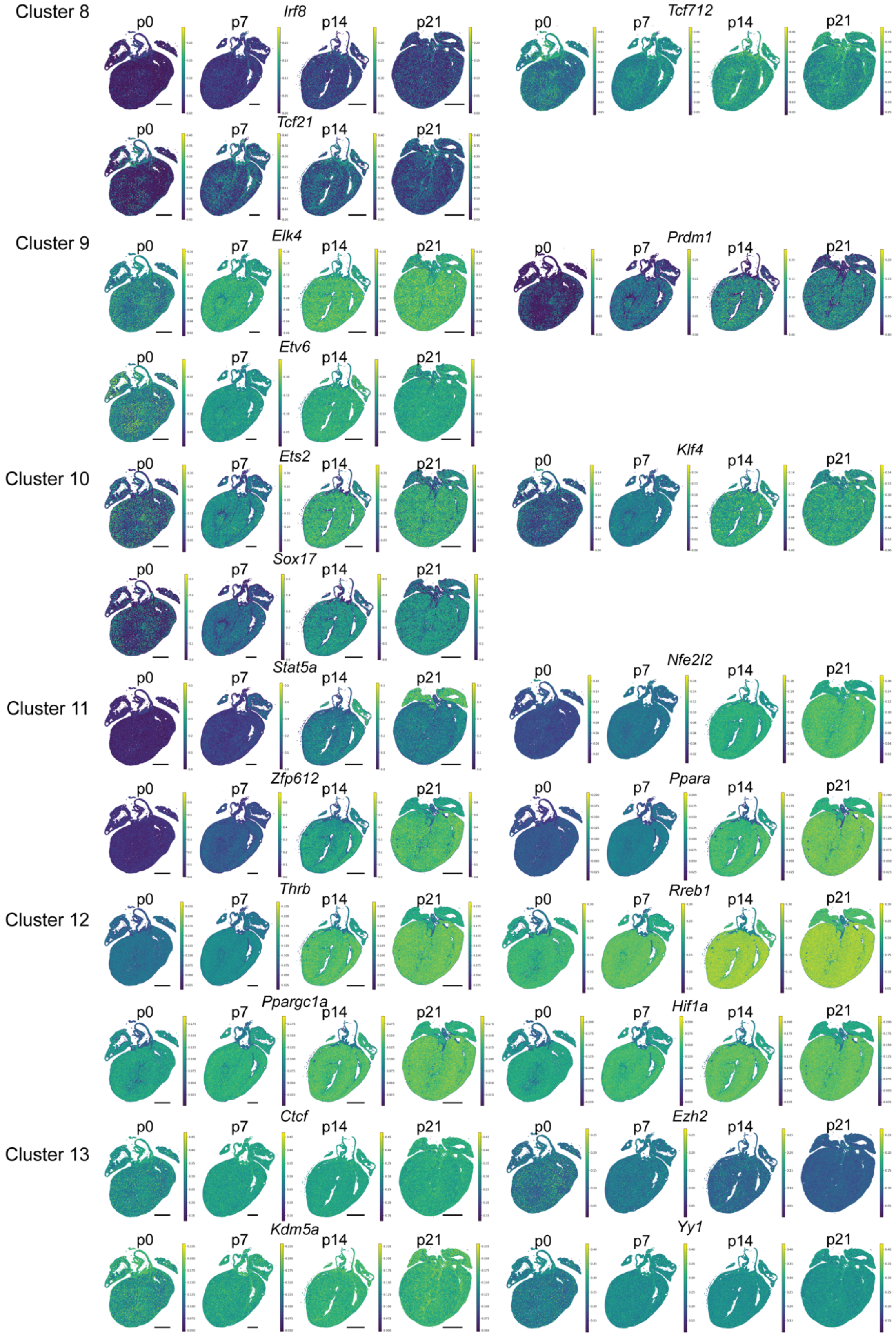


**Figure S14. Regulon activities in the postnatal development for cluster 8 to cluster 13**

Representative regulon activities were displayed in the postnatal hearts spatially and temporally. The color scheme represents the regulon activity of each cell in the postnatal heart from high (green) to low (blue). Scale bar (p0, p7), 500μm; Scale bar (p14, p21), 1000μm.

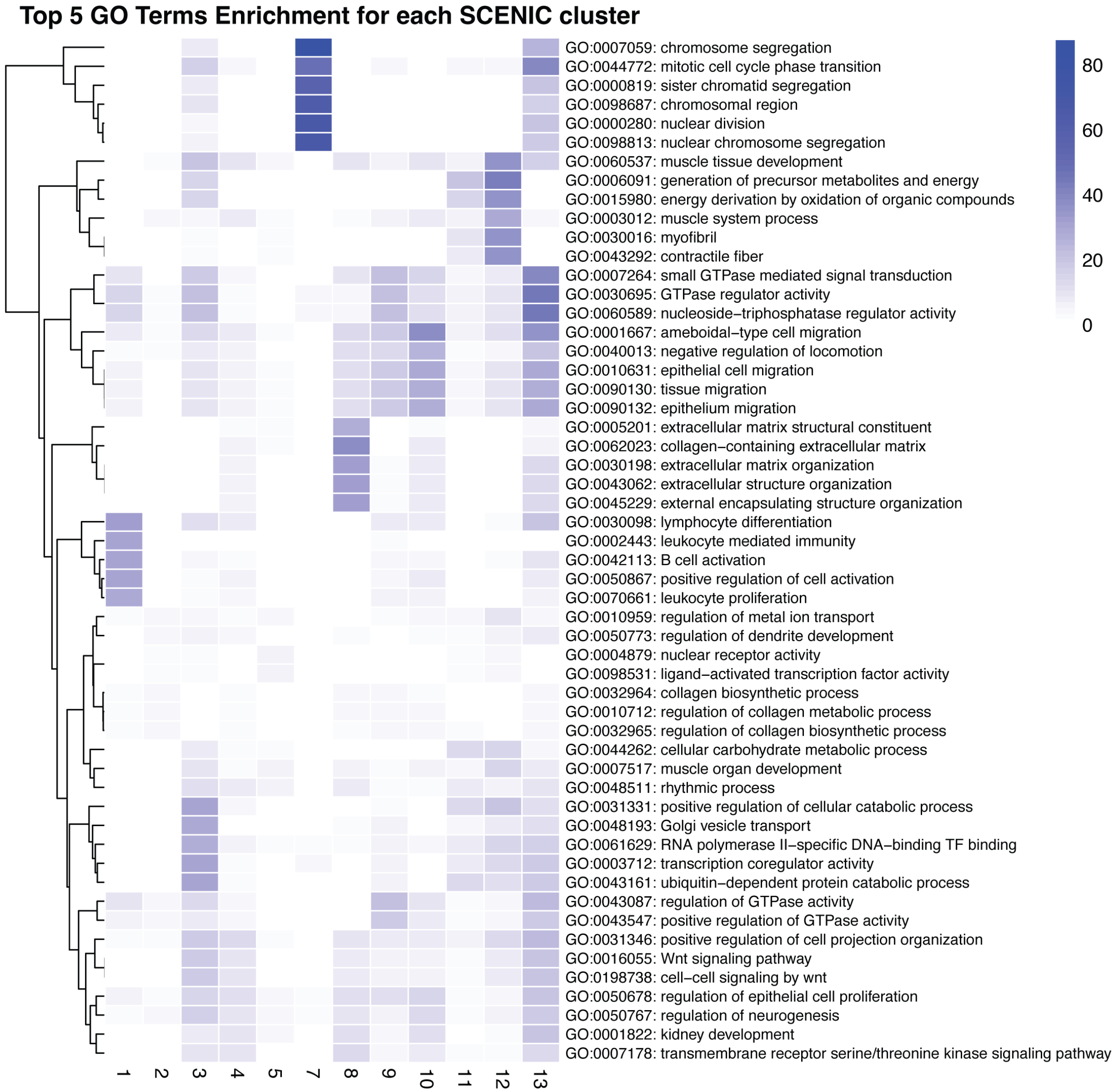


**Figure S15. Enriched gene ontology of downstream genes belong to each SCENIC cluster** Heatmap represents the top five Gene Ontology (GO) terms enriched in each SCENIC cluster. The x-axis corresponds to SCENIC clusters, while the y-axis lists the enriched GO terms. The color intensity indicates the level of enrichment, with darker shades of blue representing higher enrichment scores.

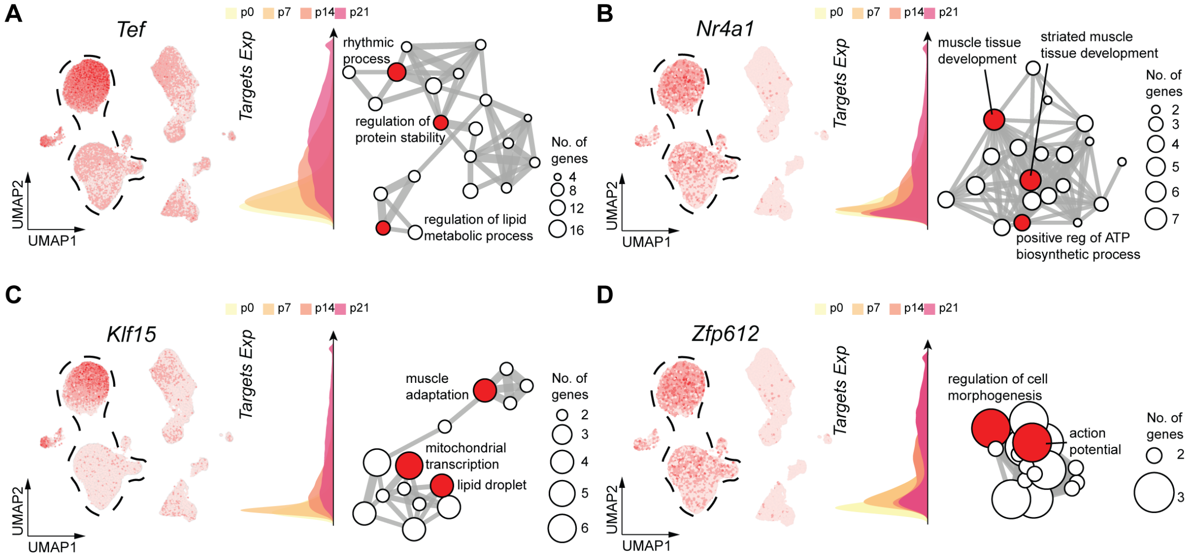


**Figure S16. Regulons' involvement in the various cardiomyocyte maturation-related programs**

A) *Tef* is a putative regulator involved in postnatal cardiomyocyte maturation. Left, the expression of *Tef* is superimposed onto the UMAP. Middle, the expression of *Tef*'s direct target at each timepoint of postnatal cardiomyocytes. Right, enriched biological process based on gene ontology enrichment analysis of *Tef*'s direct target genes.

B) *Nr4a1* is a putative regulator of postnatal cardiomyocyte maturation. Left, the expression of *Nr4a1* is superimposed onto the UMAP. Middle, the expression of *Nr4a1*'s direct target at each timepoint of postnatal cardiomyocytes. Right, enriched biological process based on gene ontology enrichment analysis of *Nr4a1*'s direct target genes.

C) *Klf15* is a putative regulator of postnatal cardiomyocyte maturation. Left, the expression of *Klf15* is superimposed onto the UMAP. Middle, the expression of *Klf15*'s direct target at each timepoint of postnatal cardiomyocytes. Right, enriched biological process based on gene ontology enrichment analysis of *Klf15*'s direct target genes.

D) *Zfp612* is a putative regulator involved in postnatal cardiomyocyte maturation. Left, the expression of *Zfp612* is superimposed onto the UMAP. Middle, the expression of *Zfp612*'s direct target at each timepoint of postnatal cardiomyocytes. Right, enriched biological process based on gene ontology enrichment analysis of *Zfp612*'s direct target genes.

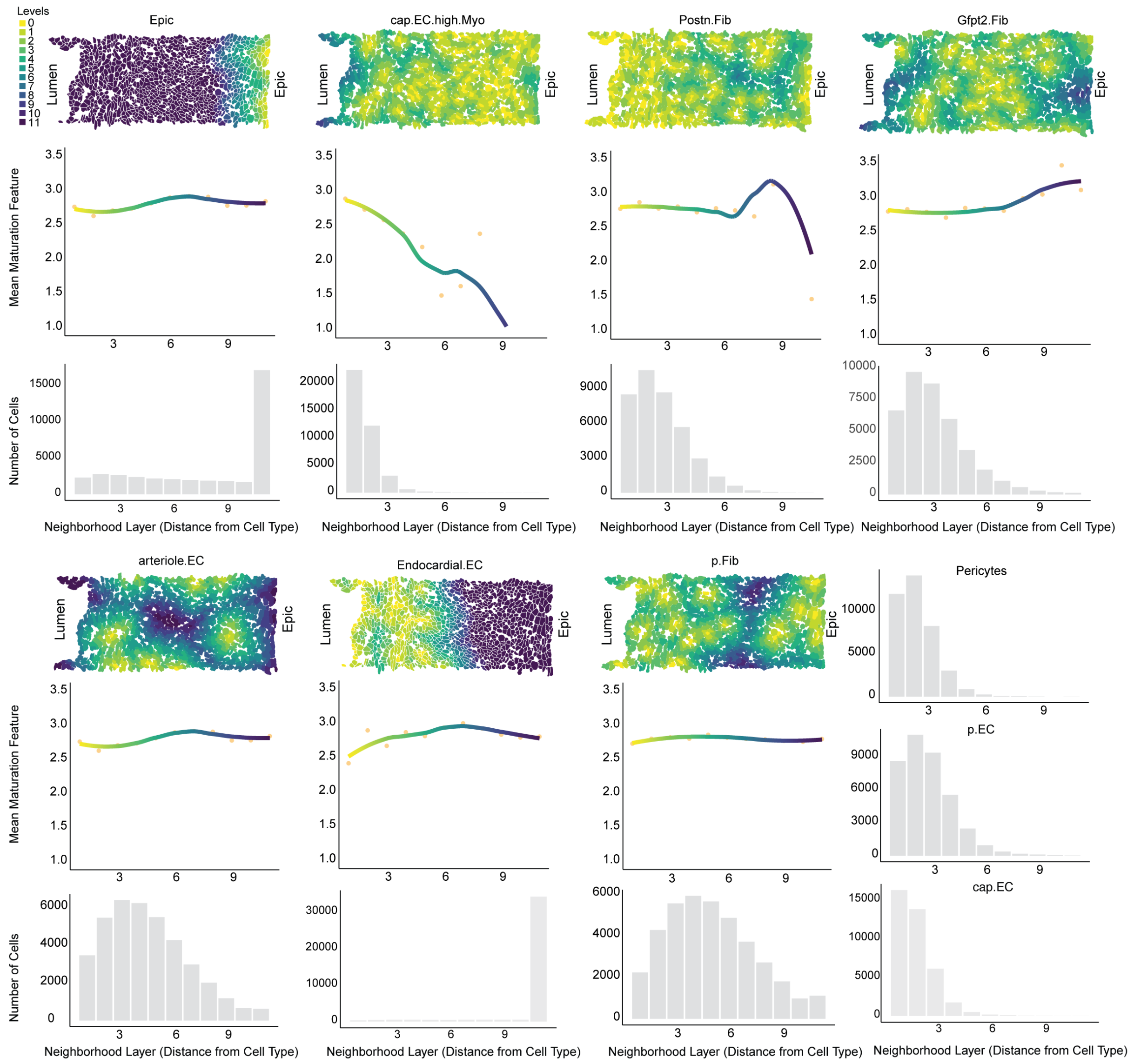


**Figure S17. Cellular neighborhoods affect the cardiomyocyte maturation**
For each non-myocyte cell type, the top panel displays the distribution of neighborhoods at a zoomed-in section of the postnatal heart at day 7. The middle panel illustrates the correlation between the maturation index and the cardiomyocyte's distance from other non-myocyte cell types. The bottom panel presents the number of cells within each distance bin corresponding to the non-myocyte cell types.

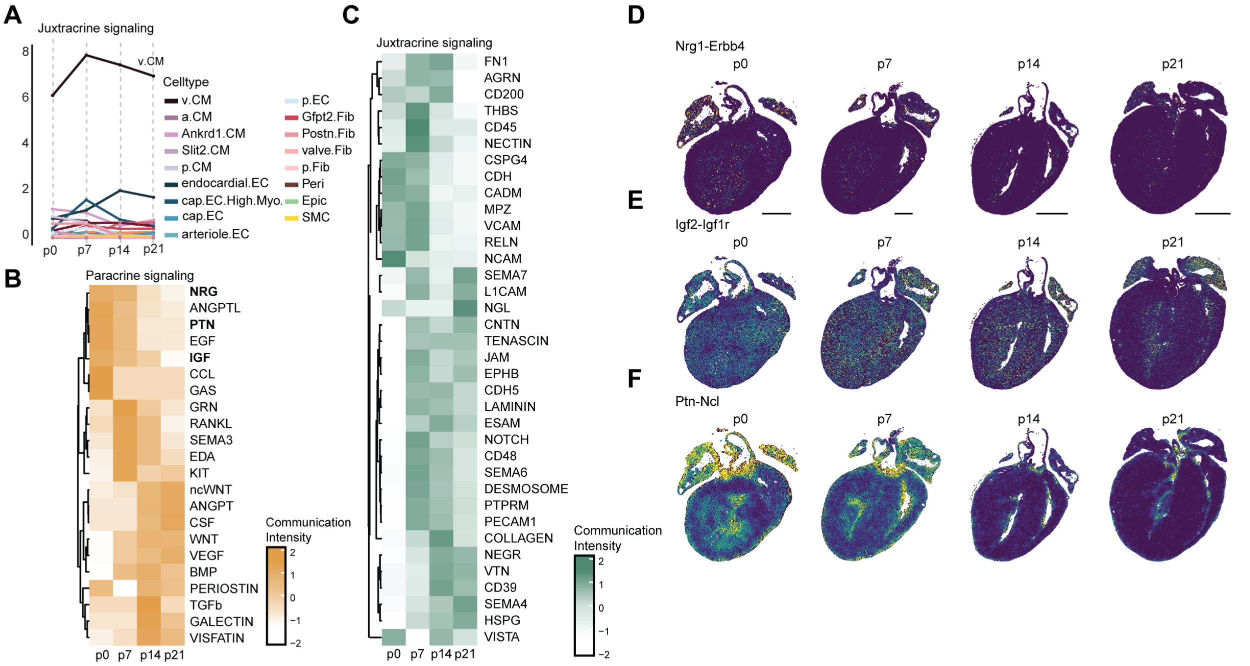


**Figure S18. Cellular communication received by ventricle cardiomyocyte during postnatal development**

A) Line plot showing the juxtracrine communication intensity received by ventricle cardiomyocytes from other cardiac cells throughout the postnatal development.

B) Heatmap showing the communication intensity of paracrine signaling pathways received by ventricle cardiomyocytes from other cardiac cells throughout postnatal development. The color scheme represents the normalized communication intensity of each signaling pathway.

C) Heatmap showing the communication intensity of juxtracrine signaling pathways received by ventricle cardiomyocytes from other cardiac cells throughout postnatal development. The color scheme represents the normalized communication intensity of each signaling pathway.

D) The spatially resolved activities of *Nrg1-Erbb4* in postnatal day 7 heart. The color Scheme represents the normalized communication intensity.

E) The spatially resolved activities of *Igf2-Igf1r* in postnatal day 7 heart. The color Scheme represents the normalized communication intensity.

F) The spatially resolved activities of *Ptn-Ncl* in postnatal day 7 heart. The color Scheme represents the normalized communication intensity.

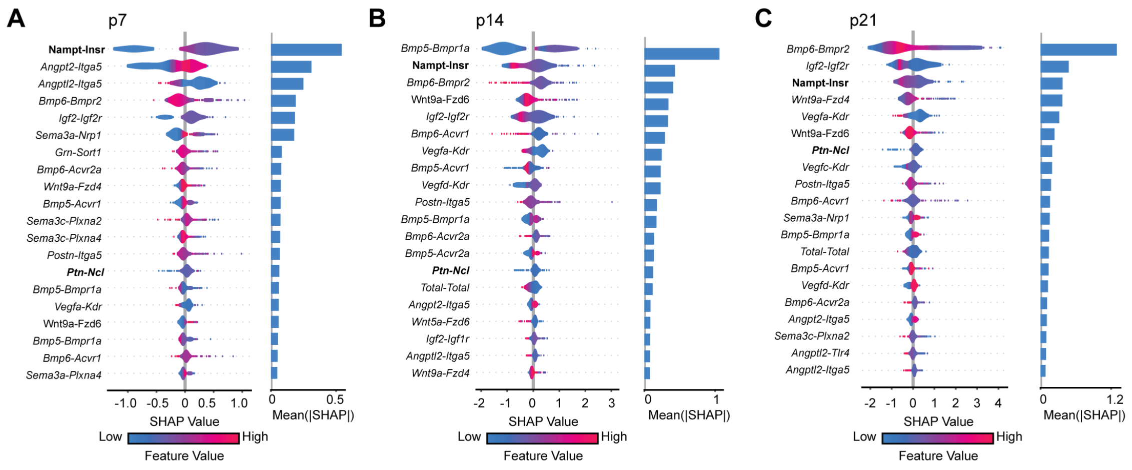


**Figure S19. Top L-R pairs contribute to the CM maturation index**

A) Left: Violin plot showing the SHAP value distribution of the L-R pairs with top 20 impact on the model output of p7 heart. Color Scheme reflects the L-R activity for each cell with red as high activity and blue as low activity. Right: Bar plot showing the average of the absolute value of SHAP value for each L-R pair.

B) Left: Violin plot showing the SHAP value distribution of the L-R pairs with top 20 impact on the model output of p14 heart. Color Scheme reflects the L-R activity for each cell with red as high activity and blue as low activity. Right: Bar plot showing the average of the absolute value of SHAP value for each L-R pair.

C) Left: Violin plot showing the SHAP value distribution of the L-R pairs with top 20 impact on the model output of p21 heart. Color Scheme reflects the L-R activity for each cell with red as high activity and blue as low activity. Right: Bar plot showing the average of the absolute value of SHAP value for each L-R pair.

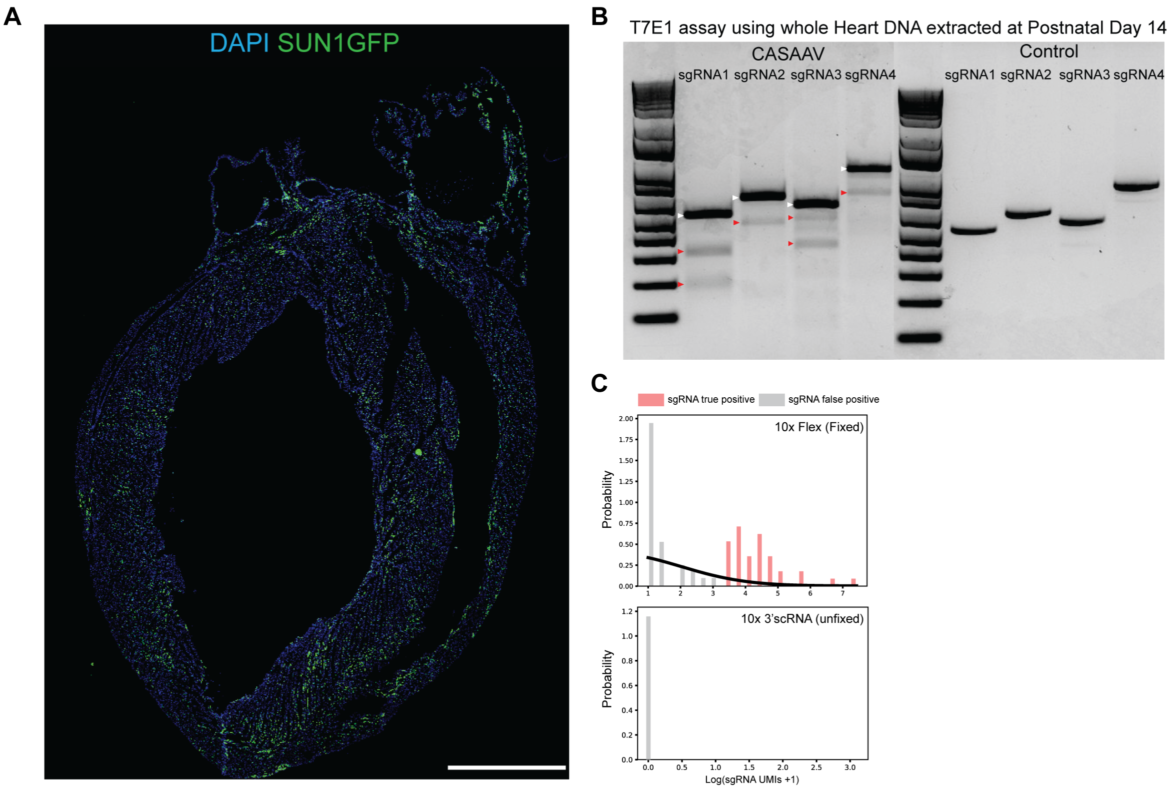


**Figure S20. Establishment of Probe-based Indel-detectable *in vivo* Perturb-seq**

A) Stitched images of a whole neonatal heart showing expression of SUN1-GFP 14 days post AAV9 injection. DAPI, nucleus, blue. SUN1-GFP, cardiomyocyte nucleus membrane, green. Scale bar, 1000 µm.

B) Agarose gel image showing the gene editing efficiency in neonatal heart with sgRNA expression.

C) Histogram showing the distribution of sgRNA reads in 10x Flex (fixed) versus 10x 3' scRNA-seq (unfixed).

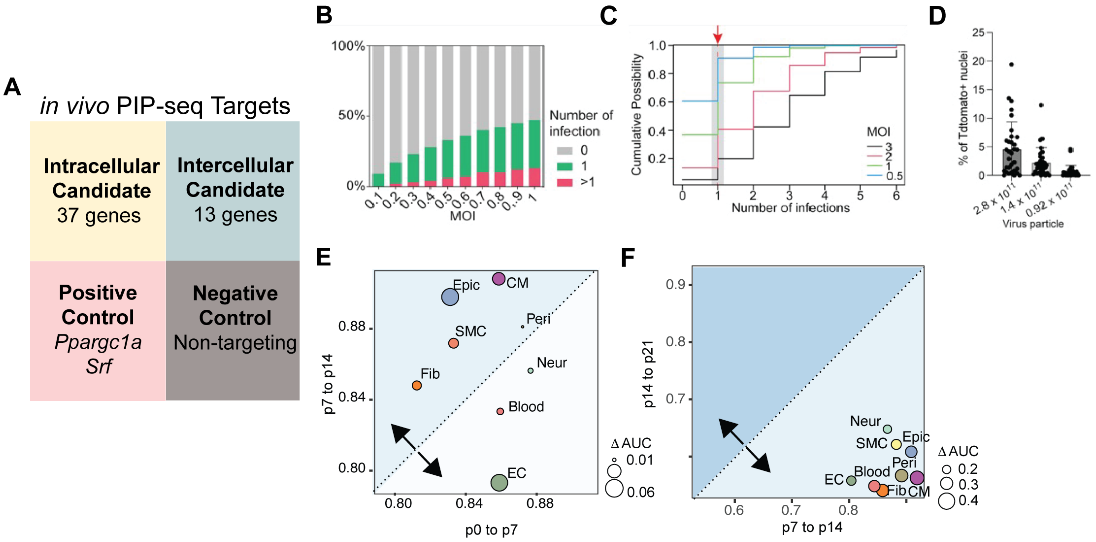


**Figure S21. Applying PIP-seq to Identify Novel Regulators of CM Maturation**

A) The library of the PIP-seq comprises 37 intracellular candidate and 13 intercellular candidate regulators that were previously identified through an integrated analysis of postnatal heart tissue. Additionally, the library includes sgRNAs targeting Ppargc1a and Srf as positive controls, while a non-targeting sgRNA serves as a negative control.

B, C) Calculated cumulative possibility of cells receiving different numbers of AAV with different MOI (0.5,1,2,3)

D) Bar plot showing the percentage of cardiomyocyte being infected with AAV with different titers of virus injected into the neonatal murine heart.

E, F) Scatter plot showing the comparison of the transcriptome dynamic between different time windows of postnatal maturation. The dot size represents the delta AUC score. Different cell types are coded as different colors of the dots.

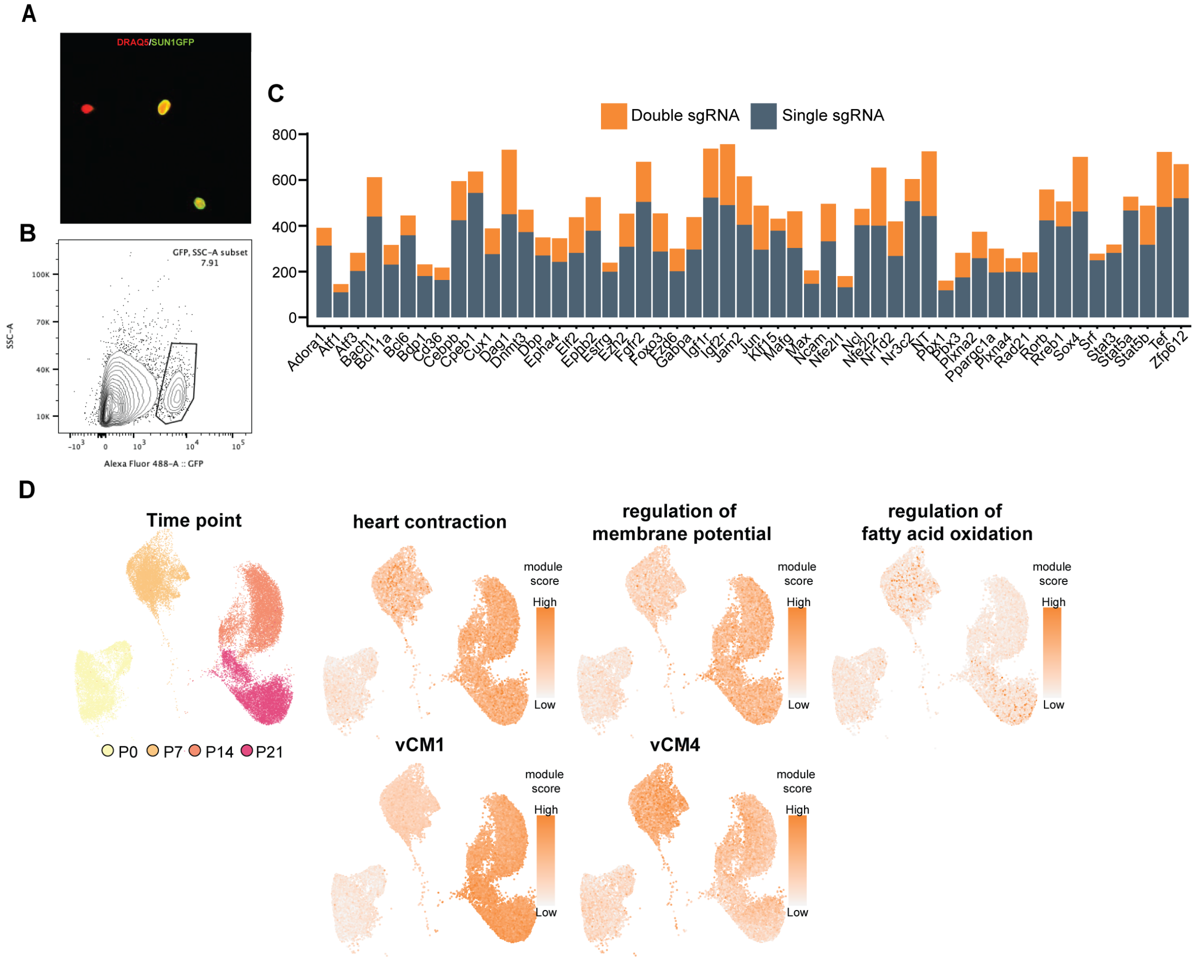


**Figure S22. Applying PIP-seq to Identify Novel Regulators of CM Maturation**

A) Immunofluorescent image showing the isolated Sun1-GFP + nuclei after flow sorting for PIP-seq preparation. Sun1-GFP, Green; DRAQ5, Red.
B) Flow cytometry plot showing the Sun1-GFP+ nuclei population takes up 7.8% of the total nuclei population, reflecting a MOI less than 0.3 for AAV9 injection in the neonatal heart.
C) Bar plot showing the number of detected nuclei harboring single (navel blue) and double sgRNAs (orange) for each perturbation target.
D) UMAP of postnatal cardiomyocyte superimposed with module score of maturation-related gene sets and WGCNA identified modules.

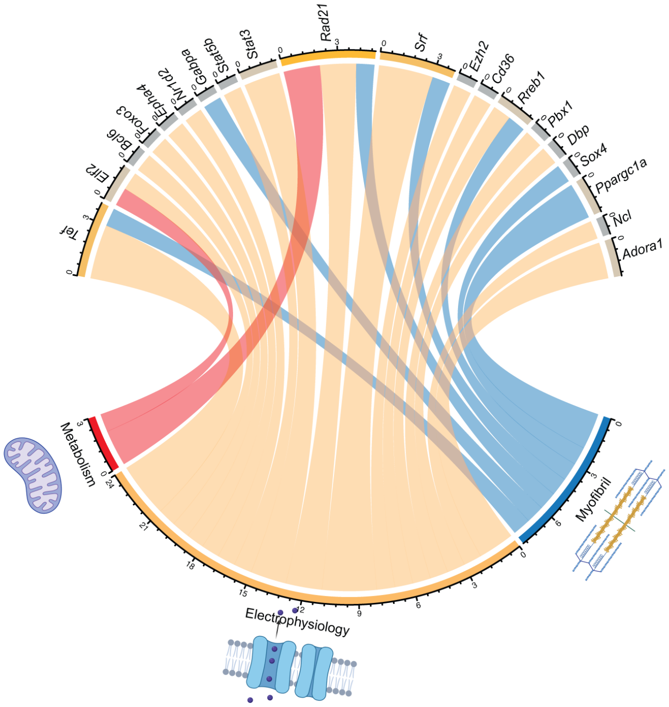


**Figure S23.** ***in vivo*** **PIP-seq identified novel regulators of postnatal cardiomyocyte maturation**
The chord diagram showing the linkage between the perturbation of candidate genes and their significant impact on the three major cardiomyocyte maturation program, Myofibril, Electrophysiology, and Metabolism.
